# Pooled screening identifies combinatorial CAR signaling domains for next-generation CAR-M immunotherapies

**DOI:** 10.1101/2025.02.16.638489

**Authors:** Yuxin Wang, Shiman Zuo, Hanjing Bao, Zehui Zhang, Yijun Chen, Wenlong Zhang, Qi Liu, Yan Lu, Yahong Huang, Wei Zheng, Nanfei Yang, Lupeng Ye, Pingping Shen

## Abstract

Chimeric antigen receptor-engineered macrophages (CAR-Ms) hold great promise for solid tumor immunotherapy. The intracellular domains (ICDs) of CARs determine the phenotypic output of therapeutic macrophages but remain largely unexplored. Here, we constructed a CAR library containing 131 unique signaling domains derived from native immune receptors and identified 17 ICDs that enhance macrophage phagocytosis, inflammatory responses, or tumor infiltration *in vitro* and *in vivo*. We further developed a scalable 3’ barcode technology, CARode, to uniquely label and trace ICD variants within large-scale combinatorial CAR library and applied it to single-cell RNA sequencing and single-cell CAR analysis to assess the synergetic effects of ICD combinations on macrophage activation. Our approach uncovered a novel CD40-LY9-FCRL1 chimeric receptor that modulates the tumor microenvironment and improves solid tumor clearance. In conclusion, our findings demonstrate that pooled screening can accelerate the discovery of complex ICD constructs, providing a powerful platform for engineering macrophage-based immunotherapies.

## Introduction

Chimeric antigen receptor T (CAR-T) cell therapy has demonstrated remarkable efficacy in the treatment of hematological malignancies [1, 2]. However, its application in solid tumors has encountered significant challenges, particularly due to limited tumor infiltration and the presence of immunosuppressive tumor microenvironment (TME) [3, 4]. Recently, CAR-engineered macrophages (CAR-Ms) were introduced as an alternative to CAR-T cell therapy for solid tumors [5, 6]. Macrophages have inherent abilities to penetrate solid tumors and modulate the TME through broad immune responses, such as phagocytosis, antigen presentation, and pro-inflammatory cytokine secretion [5, 7, 8]. To harness this potential, CAR constructs designed to enhance the phagocytic, pro-inflammatory, and tumor-infiltrating functions of macrophages are currently being investigated and have shown promising anti-tumor effects in solid tumors [9–11]. Thus, CAR-engineered macrophages with enhanced anti-tumor functions hold great potential for revolutionizing immunotherapeutic approaches for solid tumors.

The anti-tumor efficacy of CAR-Ms is strongly modulated by the intracellular domains (ICDs) of the CAR constructs [12]. Various engineering strategies have been developed for engineering the signaling domains of CAR-Ms. One current strategy involves tuning CAR signaling via the Fc receptor common gamma chain (FcRγ), which facilitates antibody-dependent cellular phagocytosis (ADCP) [13, 14]. The CD3ζ cytosolic domain, a canonical signaling domain for T cell activation, showed functional similarity with FcRγ signaling in CAR-Ms [10, 15], suggesting the compatibility of receptors that are not typically expressed by macrophages. The second strategy uses Toll-like receptor signaling to drive the M1-like phenotype of therapeutic macrophages [11, 16]. This pro-inflammatory signaling has been combined with the CD3ζ phagocytic domain in CAR-Ms to induce a bifunctional phenotype [11]. Moreover, we previously constructed a CAR bearing the CD147 signaling domain to activate the expression of matrix metalloproteinases (MMPs) to degrade the extracellular matrix (ECM) of solid tumors [9]. Taken together, different ICDs induce distinct phenotypes and functions of therapeutic macrophages. However, the optimal signaling domains required for CAR-M activation and the potential of combinatorial signaling to elicit multifunctional macrophage responses remain largely unexplored.

Advances in large-scale pooled screens, such as genome-wide CRISPR screens and gain-of-function screens, have made it possible to test gene function in a high-throughput manner [17]. Recently, this technology has been applied to CAR engineering, allowing for systematic identification of ICDs, signaling motifs, and transcription factors (TFs) that increase CAR-T cell efficacy [18–22]. However, such high-throughput approaches have yet to be applied to CAR-M optimization. Several technical challenges hinder the direct application of large-scale CAR screening to macrophage-based therapy. Unlike T cells, primary macrophages exhibit limited proliferative capacity *in vitro* [23], making it difficult to maintain and expand large cell libraries required for pooled screening. Additionally, macrophages have strong antiviral activity [24], which restricts the efficiency of viral-based CAR library delivery. The large size and complexity of CAR libraries further complicate the screening process, as each CAR variant must be effectively expressed and evaluated in parallel. Therefore, novel screening strategies designed for macrophages are required to address current limitations and advance the development of CAR constructs for macrophage-based immunotherapy.

Here, we developed a 131-member natural receptor signaling domain CAR (NaSDC) library to screen the signaling domains of immune response-related receptors as independent ICDs for CAR constructs to promote macrophage activation and anti-tumor functions. The library was screened in the human monocytic cell line THP-1 and validated in human monocyte-derived macrophages (hMDMs). We revealed that 17 ICDs have distinct anti-tumor functions, including phagocytosis, pro-inflammation, and tumor infiltration. Furthermore, we developed a large-scale combinatorial signaling domain CAR (CoSDC) library to evaluate the synergistic effects of signaling combinations of the 17 ICDs in promoting macrophage activation. The CoSDC library was constructed and barcoded via the CARode (Barcode for CAR) strategy, which facilitates unique labels of combinatorial CAR constructs in a scalable manner and is compatible with single-cell RNA sequencing (scRNA-seq). The CoSDC library screens revealed multi-CAR constructs that improved macrophage anti-tumor functions, including a CD40-LY9-FCRL1 chimeric receptor that modulated the TME and improved solid tumor clearance. Overall, these studies highlight NaSDC×CoSDC screening as a powerful method to accelerate the programming of the macrophage phenotype with enhanced anti-tumor functions.

## Results

### Design of CAR libraries for CAR-M screens

Here, we aimed to develop a platform that is suitable for high-throughput, pooled screening of CAR signaling domains within macrophages. We developed two libraries to reprogram macrophage function through single ICD CAR overexpression or combined ICD signaling (Figure 1A). The single ICD CAR library aims to assess the role of intracellular elements of natural receptors as independent signaling domains of CAR constructs in promoting macrophage functions, including phagocytosis, tumor infiltration, and pro-inflammatory responses. The combinatorial signaling domain CAR library is designed to evaluate the synergetic effects of ICD combinations on macrophage activation.

**Figure 1.**
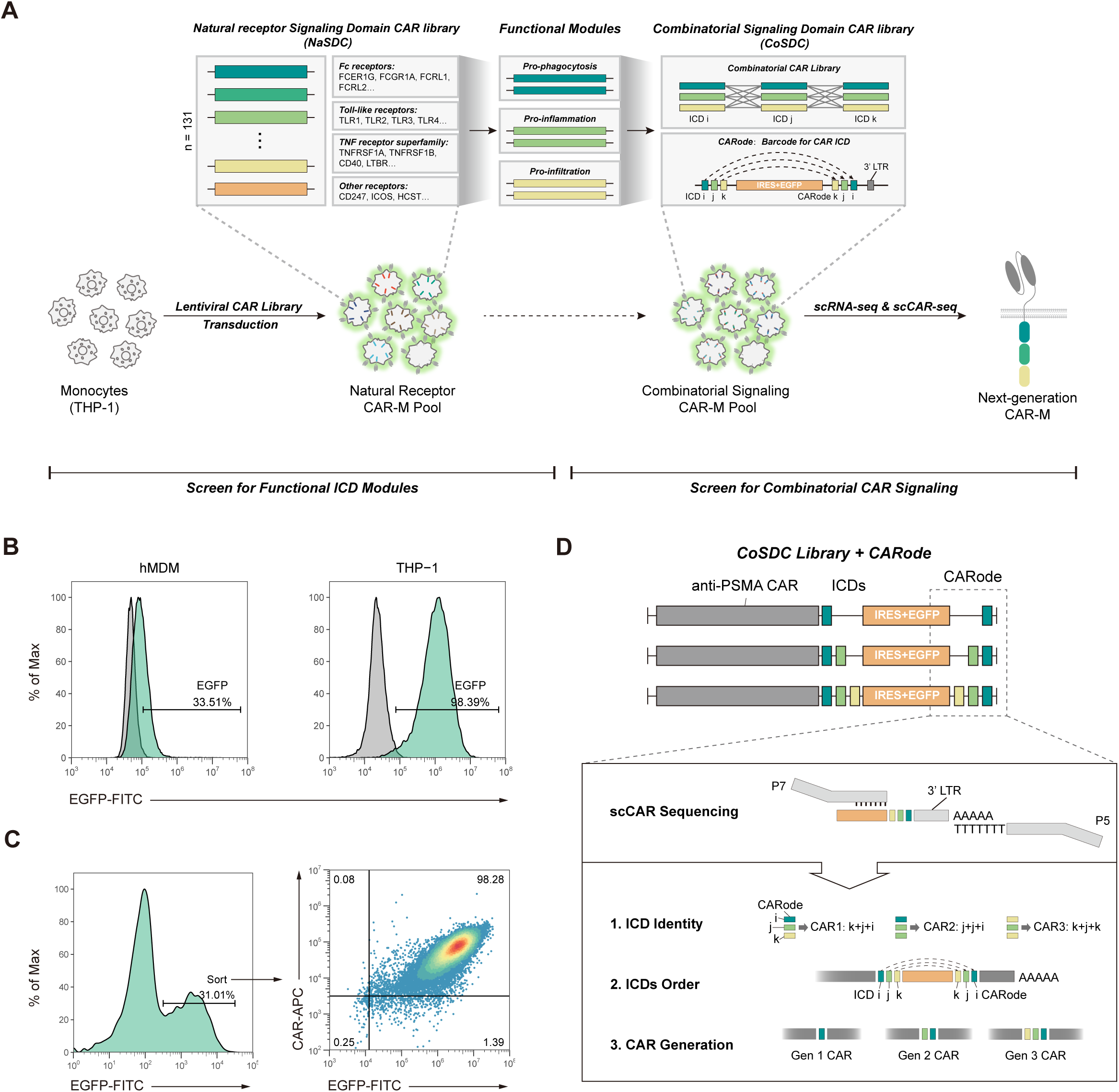
NaSDC×CoSDC screens identify therapeutic CAR-M candidates. **(A)** Schematic illustration of the NaSDC×CoSDC platform. **(B)** Flow cytometric histograms showing EGFP expression in hMDMs and THP-1 cells transduced with an EGFP reporter lentivirus. Grey peaks indicate protein expression levels in untransduced (UTD) cells. **(C)** Flow cytometric plots illustrating the NaSDC library sorting strategy (left) and CAR expression levels (right). **(D)** Schematic representation of the CoSDC library and CARode. ICD, intracellular domain. Data in (**B)** and (**C)** are representative of three independent experiments.

Human primary macrophages are resistant to virus-based gene transduction. Thus, we first needed to determine the proper cell types for large-scale CAR variant screens. The human monocytic cell line THP-1 can be differentiated into macrophages, and this approach has been used previously for genome-wide screening [25, 26]. The infectivity of lentivirus was greater in THP-1 cells than in human monocyte-derived macrophages (hMDMs) (Figure 1B). We confirmed that THP-1-derived macrophages expressed macrophage markers (Figure S1A) and showed the capacity to polarize to the M1, M2 and tumor-associated macrophage (TAM) phenotypes (Figure S1B-S1D). We then transduced CAR variants with the ICD of CD247 (CAR-3ζ) or TLR4 (CAR-t4) into THP-1 cells and observed increased phagocytosis and pro-inflammatory gene expression in THP-1-derived macrophages in response to CAR signaling (Figure S1E-S1I), suggesting that THP-1-derived macrophages constitute a proper model for CAR signaling screening studies.

To construct a library for the functional evaluation of natural receptor signaling domains as independent ICDs of CAR constructs within macrophages, we mined macrophage activation-related receptors from various protein families, such as Fc receptors, Toll-like receptors (TLRs), and the TNF receptor superfamily (Figure 1A). Given the homology of receptors across the immune system and the established role of the T cell-related receptor CD3ζ (encoded by *CD247*) as a functional cytosolic domain of CAR-Ms [10], we included receptors that are predominantly expressed in T cells, natural killer (NK) cells, dendritic cells (DCs), and B cells to determine whether subsets of these could engage in distinct and beneficial signaling in macrophages (Figure S2A). Taken together, a total of 131 signaling domains (Table S2) were collected and constructed within the context of a PSMA CAR (containing an anti-PSMA single-chain variable fragment, a CD8α hinge, and a CD8α transmembrane domain) (Figure S2B). The generated library is called the NaSDC library. A truncated CAR without a signaling domain was added to the library as a negative control. Each CAR variant was uniquely barcoded (Figure S2B) and sequenced to confirm the construct. We then generated NaSDC lentiviruses from the plasmid library to transduce THP-1 cells at a multiplicity of infection (MOI) of 0.3 to minimize multiple integrations per cell (Figure S2C). The ability of NaSDC CAR-Ms to bind to soluble PSMA antigen was confirmed by flow cytometry (Figure 1C). We observed no significant correlation between ICD length and abundance in the CAR-M library (Figure S2D). Together, the NaSDC library provides a robust platform for systematically screening diverse natural receptor signaling domains as independent ICD of CAR-Ms.

We next sought to construct a large-scale combinatorial signaling domain CAR (CoSDC) library to evaluate the synergistic effects of ICD combinations in promoting macrophage activation. A well-established method for identifying different gene variants in large-scale libraries is to use randomized 5∼20 bp barcode sequences attached to untranslated regions (UTRs) [18, 27, 28], which require pooled PCR and long-read sequencing to match the barcode sequences with gene variants. Here, we developed a novel, scalable method to barcode large-scale CAR libraries, namely CARode (Barcode for CAR). Briefly, we generated constructs with a cloning cassette that was placed between the ICD sequences and barcode sequences (Figure 1D and S3A). The cassette consisted of customized cleavage sites and a stop codon and was flanked by linkers that served as homology arms for assembly (Figure S3B). Each ICD was PCR amplified to combine with the cloning cassette and a unique 6-bp barcode and then incorporated into a PSMA CAR chassis to generate a single ICD CAR (G1) library (Figure S3A). The “2nd ICD-cassette-barcode” constructs were subsequently incorporated into the G1 library to replace the cassette of the first ICD, resulting in a second-generation (G2) CAR library with two fused barcodes (Figure S3A). The third-generation (G3) CARs were generated in a similar way (Figure S3A). This approach could theoretically enable more than 16 million possible combinatorial CARodes for the G2 library and over 68 billion CARodes for G3 CARs. Given their proximity to the 3’ UTR, these tandem CARodes allow sequencing of CAR identities in single-cell transcriptomes using commercially available 3’ capture kits (Figure 1D). Additionally, the arrangement of CARodes reflect the order of ICDs in CAR constructs, and the length of CARodes indicates the generation of CARs (Figure 1D). In summary, CoSDC screens with CARode technology enable rapid evaluation of CAR ICD combinations for engineered immunotherapies.

### NaSDC allows for the discovery of CAR-M variants with distinct anti-tumor functions

We first aimed to identify signaling domains that trigger the phagocytosis of macrophages in the context of CARs. We adopted a bead-based selection strategy for the rapid separation of CAR-Ms on the basis of their ability to phagocytose magnetic substrates (Figure 2A). After incubation with magnetic particles, NaSDC library cells were passed through a uniform magnetic field that selectively captured magnetized cells that had ingested magnetic particles, while cells that did not phagocytose beads passed through. Subsequently, RNA was isolated and reverse transcribed into cDNA, followed by barcode sequencing to compare the abundance of each CAR variant in the input and output populations.

**Figure 2.**
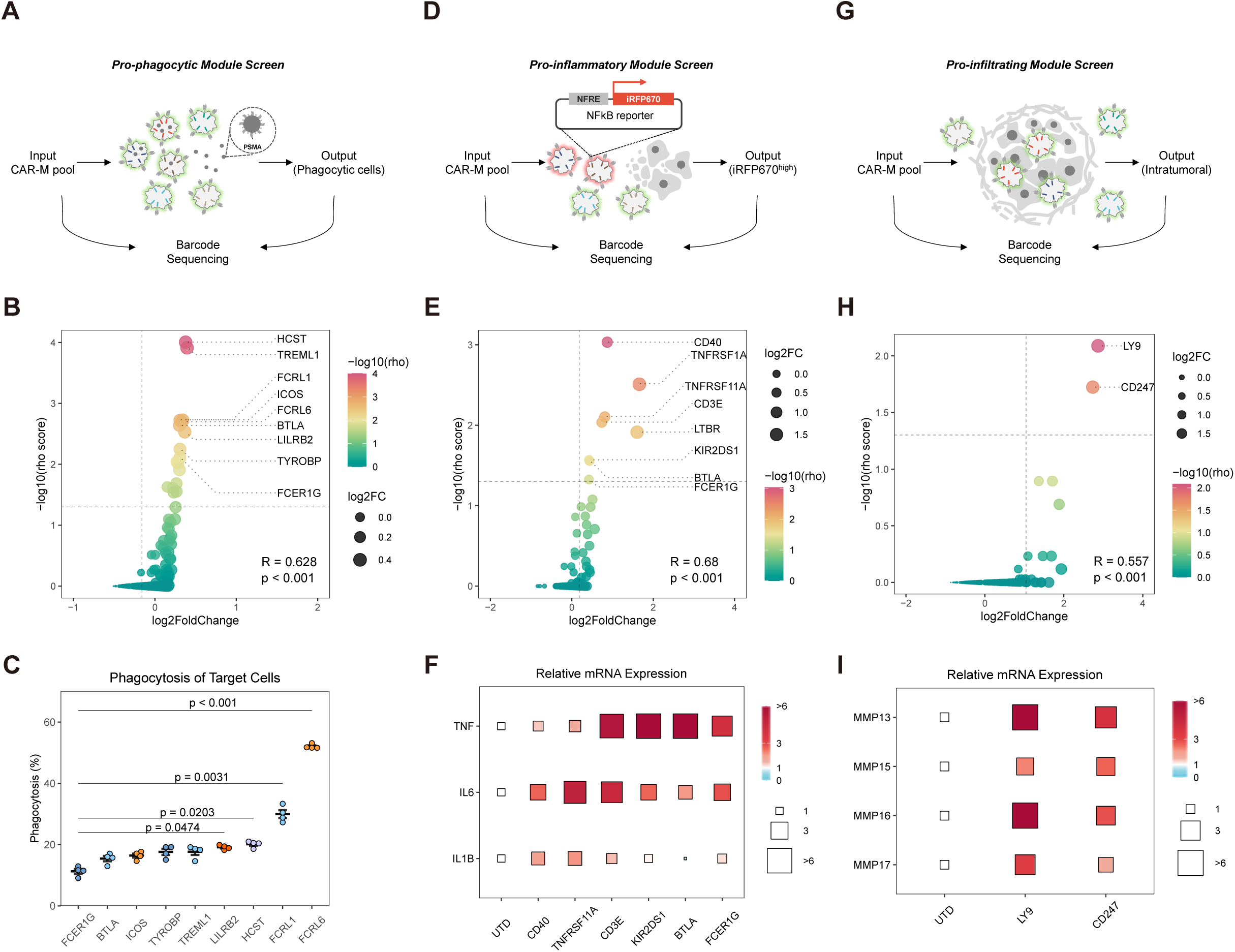
NaSDC screens reveal known and novel CAR-M candidates. **(A)** Schematic representation of the pro-phagocytic module screening strategy. **(B)** Correlation between log2 fold change and −log10 rho scores for pro-phagocytic CAR variant screens. Data represent 18 biological replicates across six NaSDC lentivirus batches. **(C)** FACS-based quantification of PC3-PSMA tumor cell phagocytosis by CAR-Ms (n = 4 biological replicates). **(D)** Schematic representation of the pro-inflammatory module screening strategy. **(E)** Correlation between log2 fold change and −log10 rho scores for pro-inflammatory CAR variant screens. Data represent 15 biological replicates across five NaSDC lentivirus batches. **(F)** Relative mRNA expression of *TNF*, *IL6* and *IL1B* in CAR-Ms cocultured with PSMA positive target tumor cells (n = 4 biological replicates). **(G)** Schematic representation of the pro-infiltrating module screening strategy. **(H)** Correlation between log2 fold change and −log10 rho scores for pro-infiltrating CAR variant screens. Data represent 9 biological replicates across three NaSDC lentivirus batches. **(I)** Relative mRNA expression of MMP family genes in CAR-Ms cocultured with PSMA positive target tumor cells (n = 4 biological replicates). Dashed lines in (**C**, **F**, **I**) indicate −log10(0.05) (horizontal) and log2 fold change of truncated CARs (CARΔ) (vertical). Data in (C) are presented as mean ± s.e.m. and analyzed using one-way ANOVA followed by Dunnett’s multiple-comparison test.

We integrated beta scores from the robust rank aggregation (RRA) model with overall fold changes to evaluate CAR enrichment in the output populations. The screening was performed in sextuplicate to enable RRA model and reduce batch effects. The results of the two algorithms were strongly correlated (Figure 2B). Notably, we found that the previously reported pro-phagocytic signaling domains FCER1G and CD247 were highly enriched in the output pools (Figure S4A), suggesting that our screening method yielded reliable results. Signaling by Fc receptor superfamily had strong effects on macrophage phagocytosis, especially FCRL1 and FCRL6, which exhibited similar pro-phagocytosis capacity compared with FCER1G signaling (Figure S4B). Among the top positive hits were HCST (encoding DAP10), which is required for NK cell activation and associated with T cell responses [29], and TREML1, which can promote the aggregation of platelets [30] (Figure 2B). Interestingly, the inhibitory signals BTLA and LILRB2, which are expressed primarily in T cells and NK cells [31, 32] (Figure 2B), also promote macrophage phagocytosis, suggesting distinct functions of these receptors in macrophages. Overall, eight signaling domains showed an enhanced ability to phagocytose magnetic substrates compared with FCER1G signaling.

We next sought to validate the pro-phagocytic function of the CAR variants in human primary macrophages. Given that macrophages are antiviral-transfected cells, we adopted a lipid nanoparticle (LNP) delivery system to transfer CAR-encoded mRNAs into primary human macrophages. LNPs exhibited high gene transduction efficiency and low phenotype perturbance in the context of primary macrophages (Figure S5A and S5B). The expression level peaked at 12 hours post-transfection and remained stable for up to 10 days in the cells (Figure S5A). We transduced the mRNAs of the pro-phagocytic CARs into hMDMs and confirmed their phagocytosis of the antigen-positive tumor cells (Figure S5D and 2C). Notably, all ICD variants exhibited higher pro-phagocytic capacity compared with CAR-FCER1G (Figure 2C). Among these variants, the CAR-FCRL6 and CAR-FCRL1 variants exhibited the best performance (Figure 2C).

Therapeutic macrophages must maintain a persistent pro-inflammatory phenotype within tumors to support their anti-tumor functions [7]. However, macrophages are plastic cells and can shift to an immunosuppressive phenotype in response to TME signals [7, 8]. To explore CAR variants that can repolarize immunosuppressive TAMs to a pro-inflammatory phenotype, we performed NFκB pathway activity screens using a fluorescent reporter (Figure 2D). The reporter was cotransduced with the NaSDC library into THP-1 cells to generate a reporter CAR-M library. We pretreated the reporter cells with conditioned media from prostate cancer cell cultures for two days to mimic the immunosuppressive TME. Downregulation of iRFP fluorescence induced by tumor media was observed (Figure S4C). The CAR-Ms were subsequently cocultured with antigen-positive target cell lines, and iRFP670^high^ cells were sorted (Figure S4D) and subjected to barcode sequencing.

Polarization screening revealed eight signaling domains with notable impacts on macrophage pro-inflammatory polarization (Figure 2E and S4E). We noted that both the TNF receptor superfamily and TLR family signals promoted NFκB activation (Figure S4F), whereas chimeric receptors with TNF superfamily intracellular domains, such as CD40, LTBR, TNFRSF1A, and TNFRSF11A, tended to perform better in this context (Figure 2E). The transduction of TNFRSF1A-CARs and LTBR-CARs into hMDMs resulted in limited surface expression and decreased cell viability even in the absence of stimulation (Figure S5C and S5E), suggesting potential stimulation-independent effects. Validation analyses revealed that the top variants can increase the pro-inflammatory phenotype and cytokine secretion (Figure 2F and S6A-S6C).

To discover CAR signaling that can promote macrophage infiltration, we next screened the NaSDC pool in a subcutaneous tumor model (Figure 2G). We subcutaneously (SC) injected PSMA-positive prostate tumors and peritumorally injected NaSDC macrophages 7 days after tumor inoculation. Four days later, the tumors were harvested, and the abundance of barcodes was measured using barcode sequencing (Figure S7A). We computed CAR variant enrichment as described above and merged the results from all of the mice. We observed high enrichment of CAR-Ms with LY9 and CD247 ICDs in the tumor tissues (Figure 2H and S4G). The potential of LY9 and CD247 signaling to increase macrophage abundance in solid tumors was validated in an independent assay in which hMDMs bearing either CAR-LY9 or CAR-3ζ were injected around the tumor sites, and both of these CAR-Ms were significantly enriched in target-positive tumors (Figure S7B). Validation analyses revealed that MMP family gene expression levels increased in these CAR-Ms during coculture with antigen-positive cells (Figure 2I, S5F, and S7C-S7E), suggesting enhanced matrix degradation capacity. We noted that these CAR variants lack the ability to induce a pro-inflammatory response in macrophages (Figure S7D and S7E). RNA-seq analysis revealed a transcriptional similarity of 46% in upregulated genes between LY9 and CD247 CAR-Ms (Figure S7F). Biological process analysis showed that CAR-LY9 and CAR-3ζ macrophages behaved more similarly in terms of cell proliferation (Figure S7G).

In summary, we identified 17 signaling domains, including two previously reported ICDs (FCER1G and CD247), that can elicit pro-phagocytic, pro-inflammatory, or pro-infiltrating functions in macrophages through NaSDC screens. These functional signaling domains hold potential for guiding the design of combinatorial signaling domain CARs that benefit to macrophage-based therapies.

### CoSDC enables pooled screening of ICD combinations via scRNA-seq and scCAR-seq

We next explored ICD combinations that could trigger combinatorial functions of macrophages via the CoSDC library. To construct the CoSDC library, we randomly integrated the 17 ICDs identified from NaSDC screens to generate second- and third-generation CAR constructs, yielding a theoretical diversity of 5219 barcoded signaling domain combinations. As described above, each signaling domain receives a unique barcode sequence to de-multiplex CAR generation, ICD type and position. Barcode sequencing revealed that the CoSDC plasmid library possessed all possible combinations, with balanced representation (Figure S8A). We then generated lentiviruses to transduce THP-1 cells at a low MOI to minimize multiple integrations per cell (Figure S8B). The CAR-M library was PMA-stimulated and cocultured with antigen-positive target cells. Macrophages bearing truncated CARs were incorporated into the pooled cocultures as negative controls. Following 24 h of coculture, CAR-Ms were isolated by FACS (Figure S8C) and processed for scRNA-seq through the 10X Genomics pipeline (Figure 3A). Barcoded CAR transcripts found in the scRNA-seq cDNA product were selectively amplified and sequenced.

**Figure 3.**
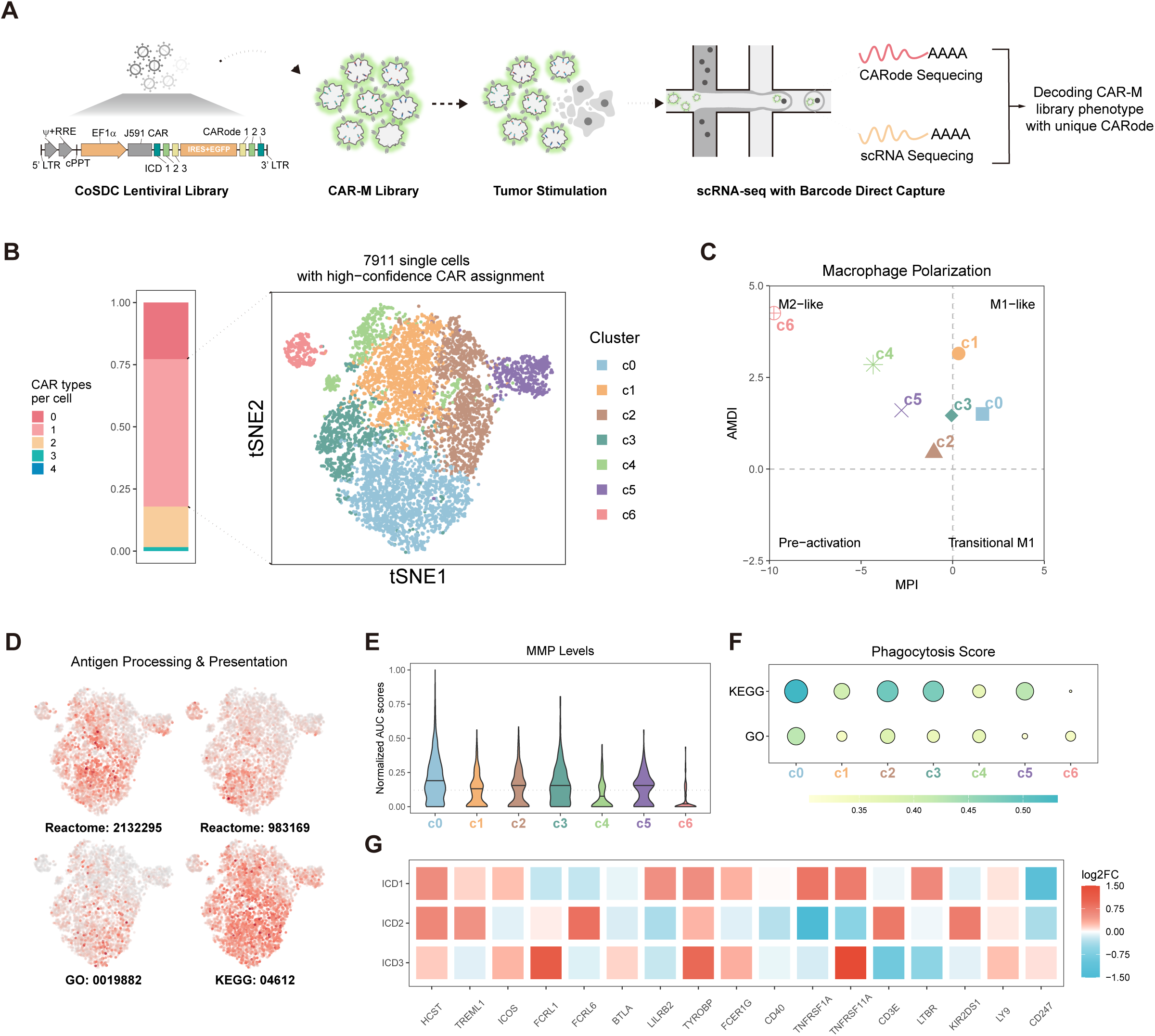
CoSDC screens identify CAR signaling combinations that enhance macrophage anti-tumor functions. **(A)** Schematic illustration of the CoSDC library and scCAR-seq screening platform. **(B)** Proportion of cells expressing different numbers of CAR variants (left), and tSNE-based dimensionality reduction and clustering (right) for single-CAR-expressing cells (right). **(C)** MacSpectrum plot showing macrophage polarization states of single-cell clusters. MPI, macrophage polarization index. AMDI, activation-induced macrophage differentiation index. Data are represented as means of clusters. Black dashed line represents the mean MPI of CARΔ. **(D)** Feature plots showing the distribution of AUC scores for antigen processing and presentation-related gene sets. Reactome gene sets “MHC class II antigen presentation” (2132295) and “Class I MHC mediated antigen processing & presentation” (983169), GO gene sets “Antigen processing and presentation” (0019882), and KEGG pathway “Antigen processing and presentation” (04612) were used. **(E)** Normalized AUC scores for MMP-related genes. Dashed line indicates the mean AUC score of CARΔ. **(F)** Dot plot showing the phagocytosis scores for single-cell clusters. KEGG pathway “Fc gamma R-mediated phagocytosis” (04666) and GO gene set “Phagocytosis” (0006909) were used. **(G)** Heatmap showing log2 fold change (c0 vs the other clusters) of each ICD in third-generation CARs.

The transcriptomes of 13,471 CAR-Ms were obtained from scRNA-seq following data preprocessing and filtering (Figure 3B). Among these cells, 58.7% could be assigned a single specific CAR variant (Figure 3B), while the remaining cells were excluded because of either the absence of CAR insertion or the presence of multiple distinct CAR constructs within the same cell. Interestingly, we noted that CAR variants triggered different functional macrophage states upon encountering target cells. Seven clusters were identified using unsupervised clustering (Figure 3B). Cluster c6 exhibited a combinatorial phenotype characterized by M2-like polarization (Figure 3C) and MHC I-mediated antigen processing and presentation (Figure 3D and S9A). Cluster c5 displayed a similar polarization pattern to that of c6 but higher levels of MMP expression (Figure 3C and 3E). Both clusters c0 and c3 significantly upregulated the expression of MMP (Figure 3E) and phagocytosis-related genes (Figure 3F), but c3 failed to transition into an M1-like phenotype (Figure 3C) and had diminished antigen processing and presentation ability (Figure 3D and S9A). Notably, CAR-Ms in cluster c0 exhibited M1-like polarization, antigen processing and presentation, MMP expression, and phagocytosis (Figure 3C-F and S9A-C), suggesting that the signaling domains within c0 CARs can trigger combinatorial anti-tumor functions of macrophages.

We next tracked the extent of enrichment for each ICD at each position in the c0 cluster. Consistent with the standard second- and third-generation of CAR-T cell constructs [1, 33], the CD247 signaling domain tended to perform better at the C-terminal end of the CAR-M constructs (Figure 3G). In contrast, some ICDs (e.g., TNFRSF1A and LTBR) are suitable for grafting into the membrane-proximal intracellular space of a CAR (Figure 3G). HCST, TYROBP, and LY9 signaling can be incorporated into any of the three intracellular positions in a third-generation CAR construct (Figure 3G). These results demonstrate the positional preferences of different ICDs when incorporated into a combinatorial signaling CAR.

### CoSDC library-guided CAR constructs induce a favorable macrophage phenotype

To validate the CoSDC-guided CAR variants and understand their effects on macrophage activation and anti-tumor functions, we selected and cloned eight CAR variants from the CoSDC library, based on the ICD features of cluster c0 (Figure 4A). Each variant consists of a pro-phagocytic module, a pro-inflammatory module, and a pro-infiltrating module. The ability of CAR-M variants to bind to PSMA antigen was confirmed by flow cytometry (Figure S10A). The ICD sizes ranged from ∼1,000–2,000 bps, and little correlation was observed between the ICD size and surface expression efficiency (Figure S10B and S10C). Interestingly, we observed significantly different CAR surface expression levels between constructs with the same ICD elements but different arrangements (COMB1: HCST-LY9-TNFRSF11A, and COMB4: TNFRSF11A-HCST-LY9) or constructs containing one different ICD (COMB2: LY9-FCER1G-CD247, and COMB3: LY9-FCER1G-KIR2DS1) (Figure S10D and S10E), suggesting that the ICD type and position modulate CAR surface expression within macrophages.

**Figure 4.**
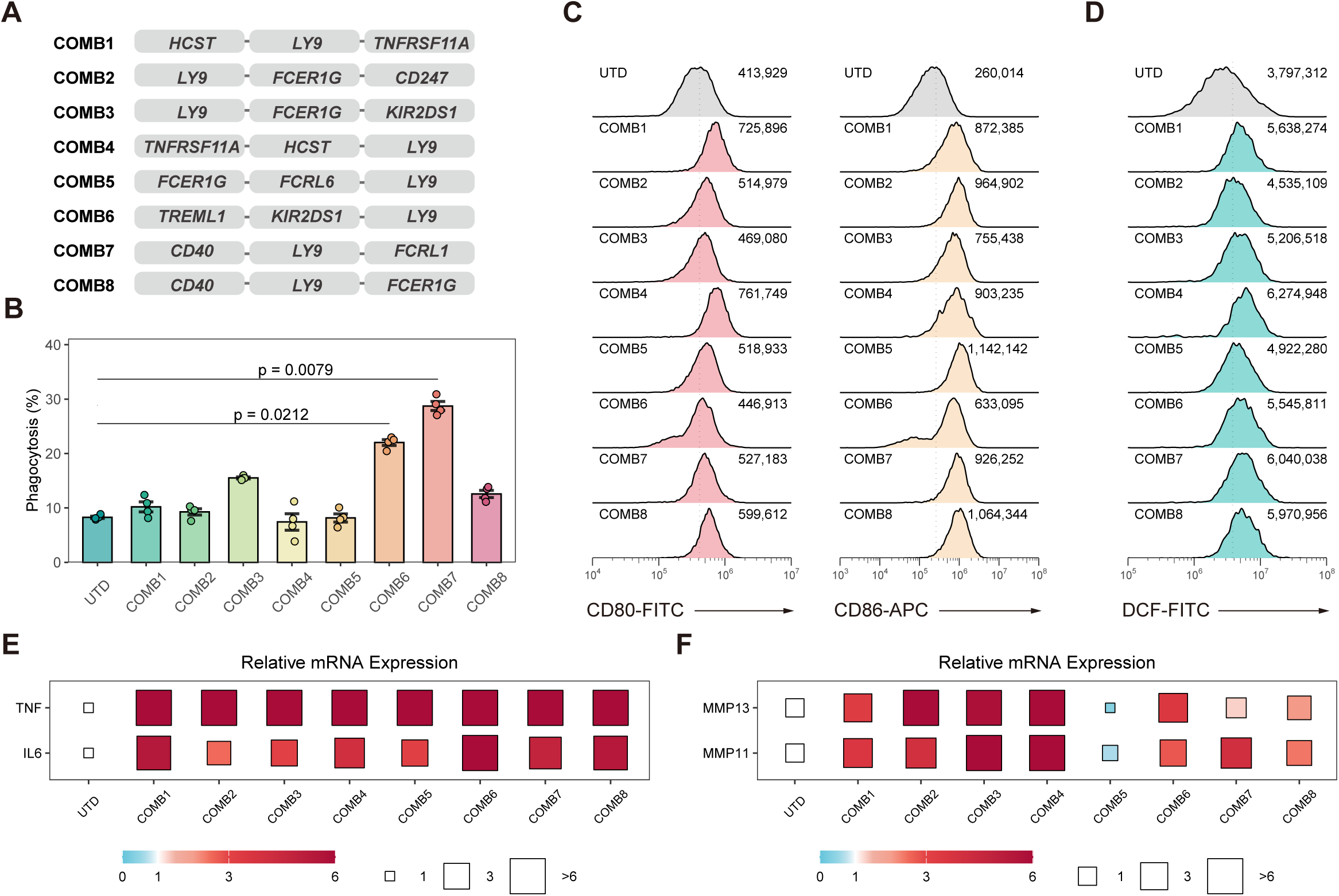
Combinatorial CAR constructs enhance macrophage phagocytosis, pro-inflammatory activation, and infiltration capacity. **(A)** Description of CAR variants used for individual characterization. **(B)** FACS-based quantification of PC3-PSMA tumor cell phagocytosis by UTD and CAR-Ms (n = 4 biological replicates). **(C)** Flow cytometric histograms showing surface expression of CD80 and CD86 in UTD and CAR-Ms cocultured with PSMA positive tumor cells. **(D)** Flow cytometric histograms of reactive oxygen species (ROS) levels in UTD and CAR-Ms cocultured with PSMA positive tumor cells. DCF, 2’-7’dichlorofluorescein. **(E)** Relative mRNA expression of *TNF* and *IL6* in UTD and CAR-Ms cocultured with PSMA positive tumor cells (n = 4 biological replicates). **(F)** Relative mRNA expression of *MMP11* and *MMP13* in UTD and CAR-Ms cocultured with PSMA positive tumor cells (n = 4 biological replicates). Data in (**B)** are presented as mean ± s.e.m. and analyzed using one-way ANOVA followed by Dunnett’s multiple-comparison. Data in (**C**, **D)** are representative of three independent experiments. Dashed lines (**C**, **D**) represent the mean fluorescence intensity (MFI) for UTD macrophages.

We next evaluated the effect of the combinatorial signaling of CARs on macrophage activation. We incubated CAR-Ms with mTagBFP2-expressing PC3-PSMA cells at an effector/target (E/T) ratio of 1/1 and observed that COMB3, COMB6, COMB7 and COMB8 CARs led to remarkable phagocytosis of tumor cells compared to UTD cells over a time course of 4 hours (Figure 4B). Among these CAR-Ms, COMB6 and COMB7 CAR-Ms showed the best performance in promoting phagocytosis, suggesting their anti-tumor activity *in vitro*. We noted that all eight CAR variants presented high CD80 and CD86 expression (Figure 4C). Consistent with these findings, the reactive oxygen species (ROS) levels (Figure 4D) and the *IL6* and *TNF* expression levels were increased in these CAR-Ms (Figure 4E), suggesting their pro-inflammatory ability. In addition, we noted that all the CAR variants except COMB5 promoted MMP family gene expression in macrophages (Figure 4F). In summary, the NaSDC and CoSDC libraries facilitated the rapid identification of CAR ICD combinations that exhibit synergetic anti-tumor effects in CAR-M immunotherapy. We chose the COMB7 CAR construct for further development due to its robust pro-phagocytic function and comparable pro-inflammatory and pro-infiltrating capabilities.

### A novel CD40-LY9-FCRL1 CAR-M modifies the TME

Given that the COMB7 construct promoted pro-inflammatory phenotypes of macrophages *in vitro,* we evaluated the interactions of COMB7 CAR-Ms within the TME in a humanized immune system mouse model (Figure 5A). Given the variability in the immune system reconstitution rates among individuals, we adopted a bilateral tumor model to ensure comparability between groups. Briefly, PSMA-positive prostate tumors were bilaterally inoculated into the subcutaneous flanks of NCG mice. After confirmation of tumor engraftment, human peripheral blood mononuclear cells (PBMCs) were transplanted to reconstitute the immune system. UTD or COMB7 CAR-Ms were injected intratumorally (IT) into either the right or left flank. Four days later, the tumors were harvested, and the CAR expression levels were measured in matched tumor tissues (Figure S11A). The human CD45-positive cells were sorted and subjected to scRNA-seq (Figure S11B). The transcriptomes of 17,432 individual cells were obtained from scRNA-seq following data filtering, of which 44.6% were macrophages and 40.4% were identified as T cells (Figure 5B and S11C).

**Figure 5.**
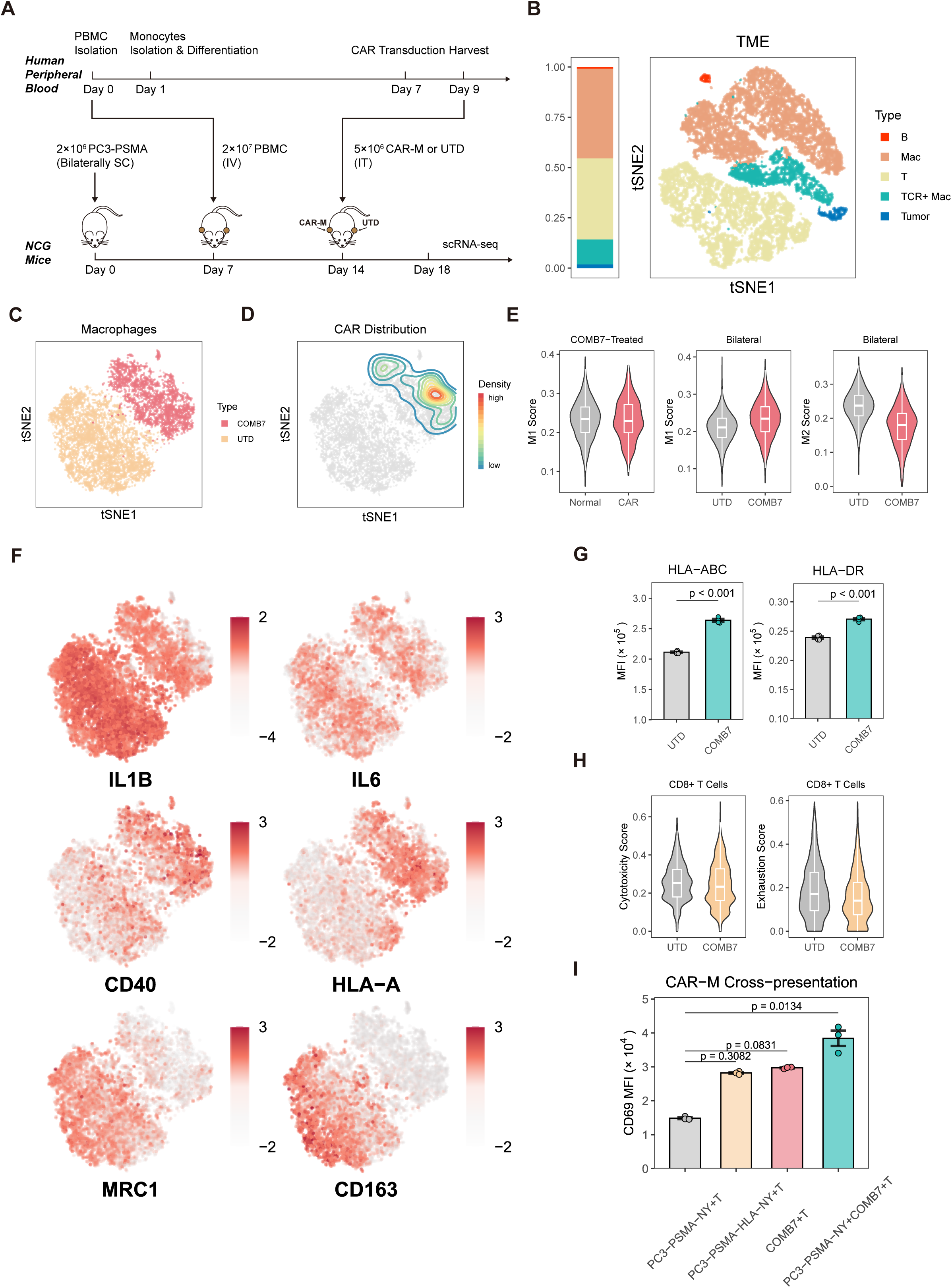
CD40-LY9-FCRL1 CAR-Ms reshape the TME. **(A)** Schematic illustration of the humanized immune system TME model. PC3-PSMA tumors were established bilaterally, followed by PBMC engraftment on day 7. On day 14, 5 × 10^6^ UTD or CAR-Ms were injected IT. Tumors were harvested on day 18 for scRNA-seq. SC, subcutaneously. IV, intravenously. IT, intratumorally. **(B)** Proportions of cell types from scRNA-seq (left) and tSNE-based clustering of all sequenced cells (right). B, B cell. T, T cell. Mac, macrophage. **(C)** Dimensionality reduction (tSNE) and clustering for macrophages. Data are colored by treatment conditions. **(D)** Density plots within tSNE maps depicting the distribution of CAR-expressing cells. **(E)** AUC scores of M1 and M2 gene sets in the indicated macrophage clusters. **(F)** Feature plots showing expression levels of selected genes. **(G)** MFI of HLA-ABC and HLA-DR in UTD and CAR-Ms after 24 h coculture with PSMA positive tumor cells (n = 3 biological replicates). **(H)** AUC scores of T cell cytotoxicity and exhaustion gene sets in the indicated T cell clusters. **(I)** CAR-M cross-presentation assay showing CD69 levels of NY-ESO-1-specific Jurkat cells (T) after coculturing with NY-ESO-1-expressing PC3-PSMA (PC3-PSMA-NY), HLA-A0201 and NY-ESO-1 co-expressing PC3-PSMA(PC3-PSMA-HLA-NY), COMB7 CAR-Ms (COMB7), and cocultures of COMB7 CAR-Ms and PC3-PSMA-NY. Unless specified otherwise, data are presented as mean ± s.e.m. and analyzed using the two-tailed Student’s t-test.

Using the scRNA-seq data, we analyzed the phenotypic and functional changes of macrophages in the TME caused by COMB7 CAR-M treatment. We isolated CAR-positive macrophages (Figure 5C and 5D) from COMB7 CAR-M-treated tumors and observed a strong M1-like polarization phenotype in these cells (Figure 5E). Notably, CAR-negative macrophages from the COMB7 flank group presented higher levels of M1 features than those from the UTD group (Figure 5E and 5F), suggesting that COMB7 CAR-Ms can maintain a pro-inflammatory microenvironment within the human TME. Although macrophages from UTD-treated tumors expressed several M1-related genes, such as *IL1B* and *IL6*, these cells simultaneously expressed the M2 markers CD163 and MRC1 (CD206) (Figure 5E and 5F). Consistent with these findings, UTD cells upregulated the expression of anti-inflammatory cytokines such as IL-10, which promote an immunosuppressive TME (Figure S11D). In addition, macrophages from the COMB7 flanks exhibited higher levels of MMP9, suggesting increased ECM degradation capacity (Figure S11D). Together, these data demonstrate that COMB7 CAR-Ms have the ability to reduce tumor-induced M2 polarization and maintain an inflammatory TME in solid tumors.

The immunosuppressive environment of solid tumors can drive exhaustion and dysfunction of T cells [34, 35]. Given the improved TME caused by COMB7 CAR-M treatment and the upregulation of costimulatory ligands as well as antigen processing and presentation genes in macrophages (Figure 5F and 5G), we examined whether COMB7 CAR-M therapy could change T cell behavior in solid tumors. We observed no significant correlations between cytotoxicity scores and COMB7 CAR-M treatment, but the exhaustion scores were notably decreased in the CD8^+^ T cells of the COMB7 group (Figure 5H). Furthermore, we tested the antigen cross-presentation ability of CAR-Ms. We cocultured the CAR-Ms with NY-ESO-1-expressing PC3-PSMA cells for 24 h and then added the Jurkat T cells expressing NY-ESO-1-specific T cell receptor (TCR) to the cocultures for 24 h. The Jurkat T cells were activated by CAR-Ms that ingested NY-ESO-1-expressing PC3-PSMA (Figure 5I). Together, these data suggest that COMB7 CAR-M treatment maintains T cell function in solid tumors.

### CD40-LY9-FCRL1 CAR improves solid tumor clearance

We next evaluated the *in vitro* killing capacity of the COMB7 CAR-induced macrophages. The COMB7 CARs were transduced to hMDMs using the LNP method. No correlation was observed between the intracellular levels of CAR-encoded mRNA and the killing efficacy (Figure S12A). We cocultured PSMA^+^ cancer cells with COMB7 CAR-Ms and observed that the COMB7 CAR improved the killing capacity across multiple E/T ratios (Figure 6A). Notably, UTD macrophages had a strong tumor killing effect at an E/T ratio of 10/1 (Figure 6A), suggesting the intrinsic anti-tumor function of primary macrophages. Compared with the CAR-3ζ, the COMB7 CAR showed increased cytotoxicity in macrophages from two human donors (Figure 6B). We next evaluated the killing capacity of the COMB7 CARs in different tumor models. We generated COMB7 CAR-Ms targeting the solid tumor antigens HER2 or GPC3 and cocultured these CAR-Ms with antigen-positive tumor cells. We observed significant killing of HER2-positive SKOV3 cells in COMB7 CAR-M cocultures (Figure 6C). Both UTD- and COMB7-engineered macrophages exhibited high killing capacity in GPC3-positive HepG2 cells (Figure 6D).

**Figure 6.**
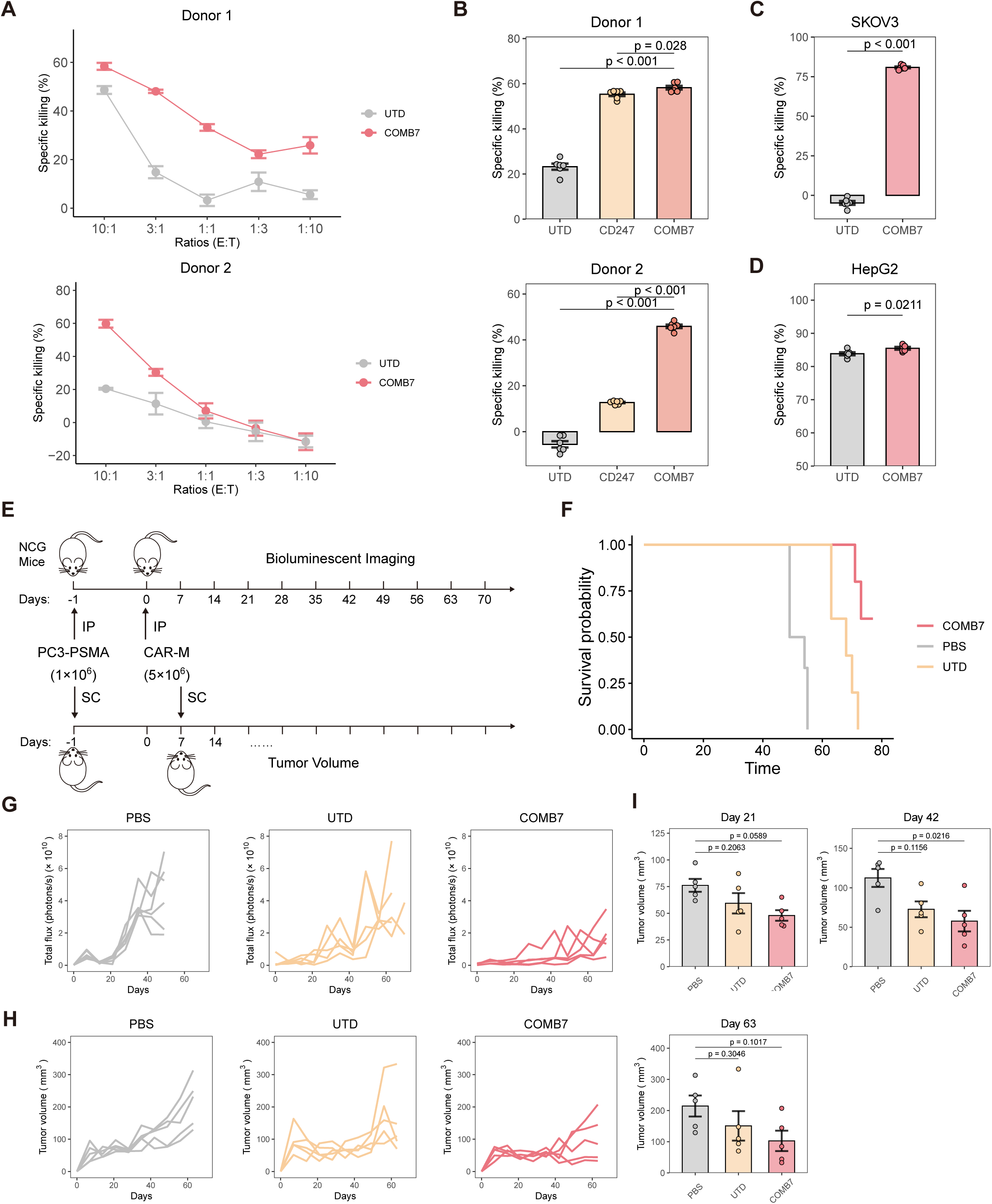
CD40-LY9-FCRL1 CAR-Ms improve solid tumor clearance *in vitro* and *in vivo*. **(A)** Specific killing of PC3-PSMA cells by UTD and COMB7 CAR-Ms at varying E/T ratios. Data represent two independent experiments from two human donors. **(B)** Specific killing of PC3-PSMA cells by UTD, CD247 CAR-Ms or COMB7 CAR-Ms at an E/T ratio of 3/1. Data collected from two human donors. **(C-D)** Specific killing of HER2^+^ SKOV3 or GPC3^+^ HepG2 cells by UTD or COMB7 CAR-Ms at an E/T ratio of 3/1 (n = 6 biological replicates). **(E)** NCG mice were IP injected with 1 × 10^6^ PC3-PSMA cells, followed by treatment with PBS, 5 × 10^6^ UTD, or 5 × 10^6^ COMB7 CAR-Ms one day later. **(F)** Overall survival for mice in each treatment group (n = 6 for PBS, n = 5 for UTD and COMB7). **(G)** Tumor burden measured by bioluminescence (total flux). Data shown are individual values (n = 6 for PBS, n = 5 for UTD and COMB7). **(H)** Tumor sizes were monitored after tumor injection. Data shown are individual values (n = 5). **(I)** Bar plot showing tumor sizes measured at 21, 42, and 63 days after tumor injection. Data are presented as mean ± s.e.m. and analyzed using one-way ANOVA followed by Dunnett’s multiple-comparison. Unless specified otherwise, data are presented as mean ± s.e.m. and analyzed using the two-tailed Student’s t-test.

We next evaluated the *in vivo* killing capacity of COMB7 CAR-Ms in NCG mice that were challenged with intraperitoneal (IP) or subcutaneous prostate tumors (Figure 6E). In the IP model, mice received a single IP administration of PBS, UTD macrophages, or anti-PSMA CAR-Ms one day post-tumor challenge. Tumor progression was monitored longitudinally using living imaging. Notably, mice treated with COMB7 CAR-Ms exhibited significantly enhanced tumor control and improved survival compared to control groups (Figure 6F, 6G and S12B). In the SC model, CAR-Ms were administered via in situ injection following confirmation of tumor engraftment (approximately 7 days post-implantation). Treatment with COMB7 CAR-Ms resulted in sustained suppression of tumor growth over a 42-day observation period (Figure 6H and 6I). Collectively, these findings demonstrate that COMB7-based chimeric antigen receptors effectively direct macrophage-mediated antitumor activity in vivo, supporting the therapeutic potential of this CAR-M platform and providing a foundation for its clinical translation.

## Discussion

The signaling domains of CAR constructs strongly modulate the phenotypic output of CAR-Ms. To discover ICDs that can enhance CAR-M anti-tumor functions, we designed a two-step NaSDC×CoSDC screening platform. The NaSDC library contains CARs with single ICDs derived from immune response-related receptors. NaSDC screening allows us to filter out ICDs with weak anti-tumor effects. CAR variants with different anti-tumor potentials were selected from NaSDC and randomly combined to generate CoSDC. We integrated CoSDC library with scRNA-seq and scCAR-seq to identify synergistic ICD combinations that enhance macrophage anti-tumor functions. As a result, NaSDC screening identified 17 functional CARs with previously unused intracellular signaling domains. Subsequent CoSDC screening using the random combinations of these 17 ICDs identified a novel CD40-LY9-FCRL1 CAR construct with enhanced anti-tumor functions. Together, the CAR candidates identified by the screening platform exhibit functions that are uncommon in current CAR-M designs and may lead to new applications for effective CAR-M therapies.

Unbiased combinatorial CAR signaling domain screens now serve as powerful tools for identifying candidates for improving T cell differentiation, activation, and cytotoxicity in CAR-T therapy [18–20, 22]. However, this screening strategy has not been applied to optimize CAR macrophages. Previous studies have focused primarily on designing CAR-Ms on the basis of known activation and polarization mechanisms of macrophages. Given the conservation of receptor signaling pathways across the immune system [36–38], we hypothesized that certain receptors, which are not naturally expressed in macrophages, could nevertheless activate these cells when incorporated into a CAR construct. Indeed, previous studies have reported T cell-related receptor CD3ζ as a phagocytic domain of CAR-Ms [10, 15]. Our NaSDC screens demonstrate that lymphatic receptors such as CD3E and ICOS can function as intracellular signaling domains to promote inflammatory responses or enhance phagocytosis in macrophages. Together, these results extend the range of signaling domains from other cell types that can act upon macrophages. Furthermore, macrophages exhibit a wide range of activation states upon sensing and responding to combined cues in their microenvironment [39–41], which cannot be accurately predicted at present. Using the CoSDC platform in combination with scRNA-seq and scCAR-seq, we found that the combinatorial signaling induced at least seven distinct macrophage activation states, although many of the CAR signaling domains have similar activation mechanisms or functions. These findings underscore the unpredictable nature of macrophage activation in response to combinatorial signaling, highlighting the critical importance of pooled screening strategies in advancing macrophage-based engineered cell therapies.

Gene transduction remains a major challenge in macrophage research. Lentiviral transduction is the primary method for large-scale library delivery. However, macrophages express retroviral restriction factors such as SAMHD1 [42–44], which limit the efficiency of lentiviral-based gene transduction. Previous pooled screening using human monocytic cell line (U937 and THP-1) to overcome these limitations [25, 26, 45]. In our study, we used PMA-differentiated THP-1 cells as an alternative model to conduct NaSDC and CoSDC screening. While this model does not fully replicate the characteristics of activated macrophages [46, 47], we observed similar phenotypic outputs between THP-1-derived macrophages and hMDMs in response to CAR signaling in NaSDC library screens, indicating its reliability for CAR identification. To enable direct screening in human primary macrophages, future research should develop more efficient gene delivery methods, such as improved viral vectors, or non-viral systems, to overcome retroviral restrictions and support CAR-M development.

Although we focused on anti-tumor functionality, NaSDC×CoSDC has potential for optimizing CAR-M therapy in various diseases, such as autoimmune diseases, cardiac injuries, and liver fibrosis. These conditions involve distinct macrophage phenotypes with diverse functions, such as anti-inflammatory responses, tissue repair, and fibrosis attenuation. Moving forward, the NaSDC×CoSDC strategy should not be limited to CAR domain screening but should be expanded to include the selection of TF combinations, cytokine combinations, and even random combinations of CARs, TFs and cytokines to further increase macrophage activation and polarization. Moreover, the integration of CoSDC with scRNA-seq can be used not only to find CAR signaling domains but also as an optimal approach for investigating the intricate mechanisms of macrophage polarization, making it possible to obtain snapshots of individual macrophages upon stimulation with different combinations of signals. Together, the NaSDC×CoSDC strategy will accelerate the process of candidate selection and design optimization for combinatorial constructs, advancing the discovery of biological mechanisms and supporting the development of a broad range of cellular therapies.

## Methods

### Cell lines

HEK293T (referred to as 293T), THP-1, PC-3, K562, Jurkat cell lines were purchased from the cell bank of the Committee on Type Culture Collection of the Chinese Academy of Sciences. 293T cells were cultured in DMEM. THP-1, PC-3 cells were maintained in RPMI 1640 medium. To obtain macrophages, THP-1 cells were differentiated with 1 ng/ml phorbol 12-myristate 13-acetate (PMA) for 48 h. PC-3 cells were transduced with a construct of FFluc-P2A-mTagBFP2 (FFLuc-BFP) to stably express firefly luciferase (FFLuc) along with a mTagBFP2 fluorescence. PC-3-FFLuc-BFP (referred to as PC3) cells were then transduced with lentiviral vectors encoding PSMA (*FOLH1*) to generate PC3-PSMA cells. K562 cells were transduced with lentiviral vectors encoding PSMA to generate K562-PSMA cells. Cells were maintained in culture medium supplemented with 10% fetal bovine serum (FBS, GIBCO) and 1% penicillin/streptomycin at 37 °C with 5% CO_2_ in a humidified incubator.

### Primary cells

De-identified healthy male and female donor blood samples were provided by the SAILYBIO Co., Ltd. Sample acquisitions were approved and regulated by the Shanghai Liquan Hospital Institutional Review Board (XF-WBC-220809). PBMCs were isolated from blood samples by density gradient centrifugation with Lymphocyte separation medium (Yeasen). Monocytes were isolated from PBMCs from all cell sources by positive selection with anti-CD14 magnetic beads (Miltenyi Biotec). For macrophage differentiation, isolated CD14^+^ monocytes were seeded on plates in X-VIVO 15 medium (Lonza) supplemented with 10% FBS and 50 ng/ml GM-CSF (Yeasen) and cultured for a further 7 days at 37 °C with 5% CO_2_.

### Xenogeneic mouse models

All mouse procedures and experiments for this study were approved by the Institutional Animal Care and Use Committee at Nanjing University (IACUC-2306005). Animals were housed in a temperature-controlled specific pathogen-free facilities with a 12 h light/dark cycle with 45-65% relative humidity. Food and water were available ad libitum. All mice were more than 8 weeks old at the beginning of the experiments. Animal sample sizes were determined based on previous studies and literatures of the field using similar experimental paradigms.

Male NOD/ShiLtJGpt-*Prkdc^em26Cd52^Il2rg^em26Cd22^*/Gpt (NCG) mice were purchased from GemPharmatech. For tumor control and survival analyses, mice were injected IP with 1×10^6^ PC3-PSMA cells. 24 hours later, 5×10^6^ CAR-Ms, UTD macrophages or PBS were injected IP. Tumor burden was monitored once per week by IP injection of D-Luciferin, Potassium Salt (Yeasen) at a dose of 150 mg/kg, then measurement of bioluminescence with an IVIS Spectrum imaging system (PerkinElmer). Tumor burden was quantified by total photon counts using the LivingImage software. For solid tumor control analyses, mice were injected SC with 1×10^6^ PC3-PSMA cells. After confirmation of tumor engraftment (about 7 days), 5×10^6^ CAR-Ms, UTD macrophages or PBS were injected IT. Tumor volume was monitored once per week. All mice were weighed weekly and were subject to routine veterinary assessment for signs of overt illness. Animals were killed at experimental termination or when predetermined IACUC rodent health endpoints were reached. For TME analyses, 4×10^6^ PC3-PSMA tumor cells were resuspended in Ceturegel Matrix LDEV-Free Matrigel (Yeasen) and injected SC into both flanks of NCG mice. After confirmation of tumor engraftment (about seven days), mice were engrafted intravenously (IV) with 2×10^7^ human PBMCs. On day 14, 5 × 10^6^ UTD or COMB7 CAR-Ms were injected IT into either the right or left flank. Four days later, the tumors were harvested for further analysis.

### NaSDC library construction

The lentiviral vector pHIV-EGFP was gifted by Bryan Welm and Zena Werb (Addgene plasmid #21373). To generate a backbone vector for ICD insertion, a codon-optimized gene encoding PSMA CAR composed of CD8α signal peptide, J591 single-chain variable fragments (scFv), CD8α hinge and transmembrane domain was commercially synthesized by GenScript and cloned into the pHIV-EGFP vector. An Esp3I cleavable linker and a randomized 18-nucleotide sequence were insert to C-terminal portion of the CAR construct. The backbone vector was digested with FastDigest Esp3I (Thermo Fisher), and 131 immune response-related ICDs (Table S2) were individually cloned into the vector using Gibson Assembly. Then, the bacterial transformation was conducted using the obtained products, and colonies were randomly selected. Sequences were performed to identify the correct encoding plasmid with a unique barcode for each CAR variant. The single CAR variant plasmids were pooled at equimolar ratios, electroporated into Endura Electrocompetent cells (LGC), and purified to generate the NaSDC plasmid library.

### Lentiviral library production

The plasmids pMD2.G and psPAX2 were gifts from Didier Trono (Addgene plasmid #12259 and #12260). To produce lentivirus, 293T cells were transfected with a plasmid library and the packaging plasmids psPAX2 and pMD2.G using Lipofectamine 2000 Transfection Reagent (Thermo Fisher) at a mass ratio of 24:3:1. Medium was changed 12 h after transfection, and the supernatant containing lentiviruses were collected in 48-72 h after transfection. The supernatant was filtered through 0.45 μm non-pyrogenic filters. To concentrate lentiviruses, the Lenti-Concentin Virus Precipitation Solution (ExCell Bio) was added to the supernatant for 12, and centrifuged for 30 min at 2,000 × *g*. The pellet was then resuspended in OptiMEM medium (Thermo Fisher) and stored at −80 °C. Medium containing concentrated lentiviral library was used to infect THP-1 cells in the presence of 8 μg/ml polybrene (Yeasen).

### Pro-phagocytic CAR variant screens

Streptavidin-coated magnetic beads (ACROBiosystems) were sterilized for 30 min in 70% isopropanol. Beads were spun down and resuspended in OptiMEM. Biotinylated human PSMA proteins (ACROBiosystems) were added to the beads at a concentration sufficient to occupy 25% of the binding sites. After incubation for 1 h, the beads were washed using PBS and resuspended in OptiMEM. The PSMA-labeled beads were added to macrophages carrying NaSDC library at a bead-to-cell ratio of 5:1. After phagocytosis for 2 h, cells were rinsed eight times by PBS to remove un-ingested particles. Then, the cells were trypsinized, harvested and sorted using a magnetic holder. The captured cells were washed three times by PBS. The input NaSDC CAR-M library and the magnetic sorted cells were store at −80 °C in TRIzol reagent (Thermo Fisher) for subsequent barcode sequencing.

### Pro-inflammatory CAR variant screens

To enable screening with fluorescent NFκB reporter system, a construct composed of NFκB response elements (NFRE), a minimal TATA-box promoter, and an iRFP670 fluorescent reporter was cloned into a pLenti6/V5 lentiviral vector to replace the CMV promotor. The fluorescent reporter was then stably co-transduced into THP-1 cells with NaSDC library, and the positive cells were selected using blasticidin (10 μg/ml). The reporter NaSDC THP-1 cells were induced by 1 ng/ml PMA for 48 h. After differentiation, PMA was washed out with PBS, and PC3 conditional media were added to the cells for 48 h to induce TAM polarization. The polarized CAR-M pool was cocultured with PC3-PSMA cells at a ratio of 1:1 for 24 h. Following coculture, cells were trypsinized, and resuspended in FACS buffer (PBS + 1-2% FBS), and iRFP670^high^ cells were sorted by a FACSAria II Cell Sorter (BD Biosciences). The input reporter NaSDC CAR-M library and the sorted cells were store at −80 °C in TRIzol reagent (Thermo Fisher) for subsequent barcode sequencing.

### Pro-infiltrating CAR variant screens

THP-1 cells were lentivirally transduced with a NaSDC library, and the positive cells were selected using flow cytometry. NCG mice were dosed with 2×10^6^ PC3-PSMA tumor cells through subcutaneous injection. The NaSDC THP-1 cells were induced by 1 ng/ml PMA for 48 h, and 1×10^7^ NaSDC CAR-Ms were peritumorally injected 7 days after tumor inoculation. Four days later, the tumors were harvested. The input NaSDC CAR-M library and the tumor tissues containing infiltrated CAR-Ms were store at −80 °C in TRIzol reagent (Thermo Fisher) for subsequent barcode sequencing.

### Barcode sequencing and data analysis

RNA was extracted from input and output population using TRIzol reagent, and reversely transcribed into cDNA using HiScript IV 1st Strand cDNA Synthesis Kit (Vazyme). Subsequently, barcode sequences were PCR amplified, and Illumina sequencing adaptors were appended using NEBNext Ultra II Q5 Master Mix (NEB). SYBR Green I (MeilunBio) was incorporated to the rection system to monitor the progression of PCR. The cDNA was amplified in a C1000 thermocycler (Bio-Rad) until the fluorescent signal approached the termination of exponential phase. Then, the products were immediately transferred to another thermocycler, and incubated at 72 °C for 5 min to complete the PCR program. The PCR products were bead-purified and sequenced by Novogene on an Illumina NovaSeq X Plus Sequencer. Considering the limited *in vitro* proliferation ability of macrophages, a single round screening would result in an elevated false positive rate within the output populations. Therefore, we repeated the screening experiments and sequencing runs to minimize the experimental bias.

DESeq2 (v1.42.1) [48] and RobustRankAggreg (v1.2.1) [49] R packages were integrated to compute the alterations of barcodes in output vs input populations. DESeq2 was used to merge the repeated experiments and calculate the overall log2 fold change (log2FC) of barcode representation. In the RobustRankAggreg analysis, the log2FC was calculated separately for each experimental batch. The barcodes were then ranked based on their log2FC and aggregated using RRA algorithm. RRA method uses rho score to retrieve the positive factors from the noise, and represent the significance probabilities for all the elements. The rho score < 0.05 and overall log2FC > log2FC (CARΔ) was considered as positive enrichment in output population.

### In vitro mRNA transcription

The genes encoding indicated CAR variants or EGFP control were codon-optimized and cloned into a plasmid carrying a T7 promoter, 5’ and 3’ UTR elements, and a poly(A) tail. The plasmids were linearized by FastDigest LguI (Thermo Fisher), and the mRNA was synthesized using T7 Co-transcription RNA Synthesis Kit (Syngene). N1-methyl-pseudouridine-5′-triphosphate (N1-Me-pUTP) was substituted for uridine triphosphate. Capping of the IVT mRNAs was performed co-transcriptionally using a trinucleotide cap1 analog CAP GAG (Syngene). The generated mRNAs were purified using VAHTS RNA Clean Beads (Vazyme). All mRNAs were analyzed by agarose gel electrophoresis and stored at −80 °C before uses. 10 pmol mRNAs were used to transduce 1×10^6^ hMDMs via Lipofectamine MessengerMAX Transfection Reagent (Thermo Fisher) according to the manufacturer’s protocol. The medium was changed 12 h after transduction, and flow cytometry was performed to detect surface expression of CAR proteins.

### Flow cytometry

To block non-specific bindings, FcR Blocking Reagent (Miltenyi Biotech) was always used for staining of monocytes, macrophages or monocytic cell lines expressing Fc receptors. Macrophage purity was tested using the following antibodies: Anti-human CD11b PE (1:100, BioLegend), Anti-human CD14 APC (1:100, BioLegend), Anti-human CD68 APC (1:100, BioLegend). M1 markers on human macrophages were detected with the following antibodies: Anti-human CD80 FITC (1:100, BioLegend), Anti-human CD86 APC (1:100, Thermo Fisher). Surface anti-PSMA CAR expression was detected using PE-Labeled Human PSMA (ACROBiosystems), or Biotinylated Human PSMA (ACROBiosystems) followed by Streptavidin/APC (Solarbio) staining. Flow cytometric analysis was performed on a NovoCyte Flow Cytometer (ACEA Biosciences) or an Attune NxT Flow Cytometer (Thermo Fisher). The data were analyzed by NovoExpress (v1.6.2) and visualized by CNSplot (v1.0.1) R package.

### Phagocytosis assay

The PC3-PSMA cells were transduced to stably express mTagBFP2 fluorescence as described above. The mTagBFP2 is a blue fluorescent protein with lower sensitivity to the acidic environment of the lysosome compared to other fluorescence proteins such as EGFP [50]. For each CAR variant, 5×10^5^ UTD or CAR-Ms were cocultured with 5×10^5^ PC3-PSMA cells for 4-6 h at 37 °C. After coculture, cells were harvested with Accutase (Yeasen) and resuspended in FACS buffer. The CAR-Ms were stained with PE-Labeled Human PSMA (ACROBiosystems), and the UTD cells were stained with Anti-human CD11b PE (BioLegend). The cells were analyzed using an Attune NxT Flow Cytometer (Thermo Fisher). The percent of mTagBFP2^+^ events within the PE^+^ population was plotted as percentage phagocytosis.

### Quantitative real-time PCR (qPCR)

Total RNA was isolated from cells or tumor tissue samples with TRIzol reagent. Reverse transcription was performed with HiScript III RT SuperMix for qPCR (Vazyme). The qPCR assays were performed on a QuantStudioLJ3 Real-Time PCR Systems (Thermo Fisher) using AceQ qPCR SYBR Green Master Mix (Vazyme) according to the manufacturer’s protocol. The data were analyzed by QuantStudio (v1.5.2) and visualized by CNSplot (v1.0.1) R package. The expression of genes was normalized to housekeeping gene *GAPDH*, and data were normalized to the mean of the control group. Primer sequences are described in Table S1.

### Cytokine assays

For each CAR variant, 1×10^5^ UTD or CAR-Ms were cocultured with antigen-positive tumor cells in an E/T ratio of 1/1 in complete X-VIVO15 medium and incubated at 37 °C for approximately 24 h. After stimulation, the supernatants were collected and IL-1β, IL-6 or TNFα were measured by ELISA following the manufacturer’s protocol (Thermo Fisher).

### RNA sequencing and data analysis

For bulk RNA-sequencing, K562 and K562-PSMA cells were labeled with green fluorescence using carboxyfluorescein succinimidyl ester (CFSE, Yeasen) method. The hMDMs from independent healthy donors were transduced with the mRNA of indicated CAR variants, and cocultured with either K562 or K562-PSMA cells at a E/T ratio of 1/1 for 24 h. Subsequently, the tumor cells were removed from the cocultures. The absolute removal of tumor cells was confirmed by flow cytometry. Total RNA was extracted from 5×10^6^ CAR-Ms using TRIzol reagent according to the manufacturer’s protocol. Next generation sequencing library preparations were conducted according to the manufacturer’s protocol. The libraries were sequenced by Novogene on an Illumina NovaSeq X Plus Sequencer. Read quality was assessed with FastQC (v0.12.1) and adapters were trimmed using Cutadapt (v4.4). Reads were then aligned to the GRCh38 genome using HISAT2 (v2.2.1) [51] and reads in exons were counted using featureCounts function in Subread (v2.0.3) package [52]. Analysis of differentially expressed genes was conducted using DESeq2 (v1.40.1) R package. Go analysis was performed using clusterProfiler (v4.10.1) R package [53]. Count data was transformed to log2 scale and normalized with respect to library size using rlog transformation produces. Absolute log2 fold change (log2FC) > 1 and p value < 0.05 was considered statistically significant.

### CoSDC library construction using CARode method

The lentiviral vector pLenti6/V5 was used as the backbone vector for CoSDC library construction, because it uses a shorter 3’ LTR region, which is suitable for 3’ capture kits of scRNA-seq. A codon-optimized gene encoding PSMA CAR composed of CD8α signal peptide, J591 single-chain variable fragments (scFv), CD8α hinge and transmembrane domain was commercially synthesized by GenScript and cloned into the pLenti6/V5 vector. A linker containing XbaI and XhoI restriction sites was insert to C-terminal portion of the CAR construct. ICDs were PCR amplified to combine with first round linkers, a cloning cassette (containing BamHI and XhoI restriction sites) and a unique 6-bp barcode (a linker-cassette-barcode-linker construct). The PSMA CAR vector was digested with FastDigest XbaI (Thermo Fisher) and FastDigest XhoI (Thermo Fisher). The ICDs were individually cloned into the PSMA CAR vector using Gibson Assembly. Then, the bacterial transformation was conducted using the obtained products. The bacteria containing different CAR variant vectors were pooled at equal OD ratios, and lysed to extract G1 library plasmid. The single ICD fragments were PCR amplified again to combine with second round “linker-cassette-barcode-linker”. The G1 library was digested with FastDigest BamHI (Thermo Fisher) and FastDigest XhoI (Thermo Fisher). The 2nd ICDs were individually cloned into the G1 to generate G2 library. Similarly, ICDs were PCR amplified to combine with third round “linker-cassette-barcode-linker”. The G2 library was digested with BamHI and XhoI enzyme. The 3rd ICDs were individually cloned into the G2 to generate G3 library. The G1, G2 and G3 library and a truncated CAR were pooled together, and digested with FastDigest BcuI (Thermo Fisher). The internal ribosome entry site (IRES) elements and EGFP fluorescence were inserted into the cloning cassette. The products were electroporated into Endura Electrocompetent cells (LGC), and purified to generate the CoSDC plasmid library.

### Single-cell RNA sequencing and single-cell CAR sequencing

The CoSDC lentiviral library was generated as described above, and transduced to THP-1 cells at a MOI of 0.3 to minimize multiple integrations per cell. The EGFP^+^ cells were sorted, PMA-induced and cocultured with high PSMA-expressing tumor cell PC3-PSMA at an E/T ratio of 1/1 for 24 h. Immediately after coculture, EGFP^+^ CAR-Ms were sorted by FACS, and scRNA-seq library was constructed using the 10X Genomics Chromium system following manufacturer’s instructions. In brief, the suspended cells were loaded into Chromium microfluidic chips and barcoded with a 10× Chromium Controller (10X Genomics). RNA from the barcoded cells was subsequently reverse-transcribed. The barcoded cDNA was purified and amplified, followed by fragmenting, A-tailing and ligation with adaptors. The 10X scRNA-seq libraries were sequenced by Novogene on an Illumina NovaSeq X Plus Sequencer. For scCAR-seq, CARode sequences were PCR amplified from 100 ng unfragmented 10X cDNA, and Illumina sequencing adaptors were appended using NEBNext Ultra II Q5 Master Mix (NEB). The CARode libraries were sequenced by Novogene on an Illumina NovaSeq X Plus Sequencer.

For scRNA-seq analysis, raw 10X sequencing reads were aligned to the GRCh38 genome using Cell Ranger (v7.2.0). Gene expression profiles were processed using Seurat (v5.1.0) R package [54]. Briefly, quality control was applied to cells based on the total UMI counts and number of detected genes. Cells with less than 5,000 UMI count were filtered, as well as cells with more than 10% mitochondrial gene count. To remove potential doublets, cells with UMI count above 70,000 and detected genes above 9,000 are filtered out. The gene counts were normalized using SCTransform. For clustering, top 2,000 variable genes were used, and ribosomal and mitochondrial genes were removed from the variable gene set before running PCA and identifying clusters using a shared nearest neighbor (SNN) modularity optimization-based clustering algorithm. The Seurat normalized count matrix was used for the downstream analysis.

For scCAR-seq analysis, the CARode sequences for each cell were extracted from the raw sequencing reads and mapped to signal domains. A total of over 21 million CARode reads were obtained from 16,612 single cells. Cells with more than 10 CARode reads were classified as CAR-Ms. 7911 cells expressing a single specific CAR variant were identified from the scRNA-seq data for subsequent clustering and functional analyses.

### Gene set activity analysis

AUCell (v1.24.0) R package [55] was employed to assess the gene set activities in scRNA-seq data. AUCell uses the “Area Under the Curve” (AUC) score to quantify the expression level of an input gene set for each cell. Previously established M1 and M2 signatures [56] were used for macrophage polarization analysis. MacSpectrum (v0.1.0) R package [57] was also employed to assess macrophage activation. To evaluate antigen processing and presentation, gene sets from the Reactome (2132295, 983169), KEGG (04612), and GO (0019882) databases were used. Additionally, KEGG (04666) and GO (0006909) gene sets were used to estimate phagocytic activity. MMP family genes MMP1-28 were applied to analyze MMP expression levels.

### Antigen cross-presentation assay

PC3-PSMA cells were transduced with lentivirus encoding NY-ESO-1 or HLA-A0201-P2A-NY-ESO-1 to generate PC3-PSMA-NY and PC3-PSMA-HLA-NY cells. NY-ESO-1-specific T cells were generated by transducing Jurkat cells with lentivirus encoding the 1G4 anti-NYESO-1 specific TCR. CAR-Ms were generated from HLA-A0201^+^ healthy donors. CAR-Ms were cocultured with PC3-PSMA-NY cells at an E/T ratio of 1/1. Subsequently, Jurkat cells were added to the cocultures at an Jurkat/CAR-M ratio of 1/1. Twenty-four hours later, Jurkat cells were analyzed for activation markers (CD69) by FACS.

### Cytotoxicity assay

To measure specific killing, a bioluminescence (BLI)-based cytotoxicity assay was performed. FFluc-positive PC3-PSMA, SKOV3, and HepG2 tumor cells were used as targets. CAR-Ms were cocultured with 10,000 target cells at the specified E/T ratios in complete X-VIVO15 medium on black 96-well flat-bottom plates. The BLI was measured using a luminometer (Tecan Spark) in relative light units (RLU). A spontaneous death control was used including tumor cells alone. The maximal death control contained 1% Triton X-100 and tumor cells. Specific lysis was calculated using: %Specific Lysis = 100 × (average spontaneous death RLU − test RLU)/(average spontaneous death RLU − average maximal death RLU). Negative specific lysis values indicate more signal than in the spontaneous death control wells.

### Statistical analysis

Statistical analysis was performed using rstatix (v0.7.2) R package. Statistical details for all experiments can be found in the figure legends. Error bars indicated the Standard Error of Mean (s.e.m.). Differences were analyzed by unpaired Student’s t test or one-way ANOVA depending on experimental conditions. Unless otherwise indicated, the experiments were performed independently in triplicate, and n is indicated in the figure legends.

## Acknowledgements

Thanks to members of Shen’s laboratory for helpful discussions, technical assistance, and critical reading of the manuscript. We thank Fuchou Tang and Lu Wen from Peking University, and Jiaming Su from Zhejiang University for their help in library construction, high-throughput sequencing and lentiviral transduction. This work was supported by grants from the Natural Science Foundation of Jiangsu Province (BK20222009 to P. Shen), the Pivotal Life Sciences Sponsorship Program (1352806 to P. Shen), the Program from National Center of Technology Innovation for Biopharmaceuticals (NCTIB2023XB01013 to P. Shen), the Program from Proof of Concept Center of Jiangsu Province (Research, Development and Clinical Study of Chimeric Antigen Receptor Macrophage Immunotherapy Technology Targeting Malignant Prostate Cancer), and the Guangdong Basic and Applied Basic Research Foundation (2021B1515120016 to P. Shen).

## Author contributions

Y. Wang and S. Zuo conceived the project and performed experiments and data analysis. H. Bao, Z. Zhang, and Q. Liu performed experiments. Y. Chen, W. Zhang, Y. Lu, Y. Huang, W. Zheng, N. Yang, and L. Ye provided scientific suggestions and contributed to the manuscript revision. P. Shen supervised the project and wrote the manuscript. The author(s) read and approved the final manuscript.

## Data availability

RNA-seq, scRNA-seq, scCAR-seq data have been deposited in the Gene Expression Omnibus repository under accession code GSE289202.

