## Supplementary figures for "Pooled screening identifies combinatorial CAR signaling domains for next-generation CAR-M immunotherapies"

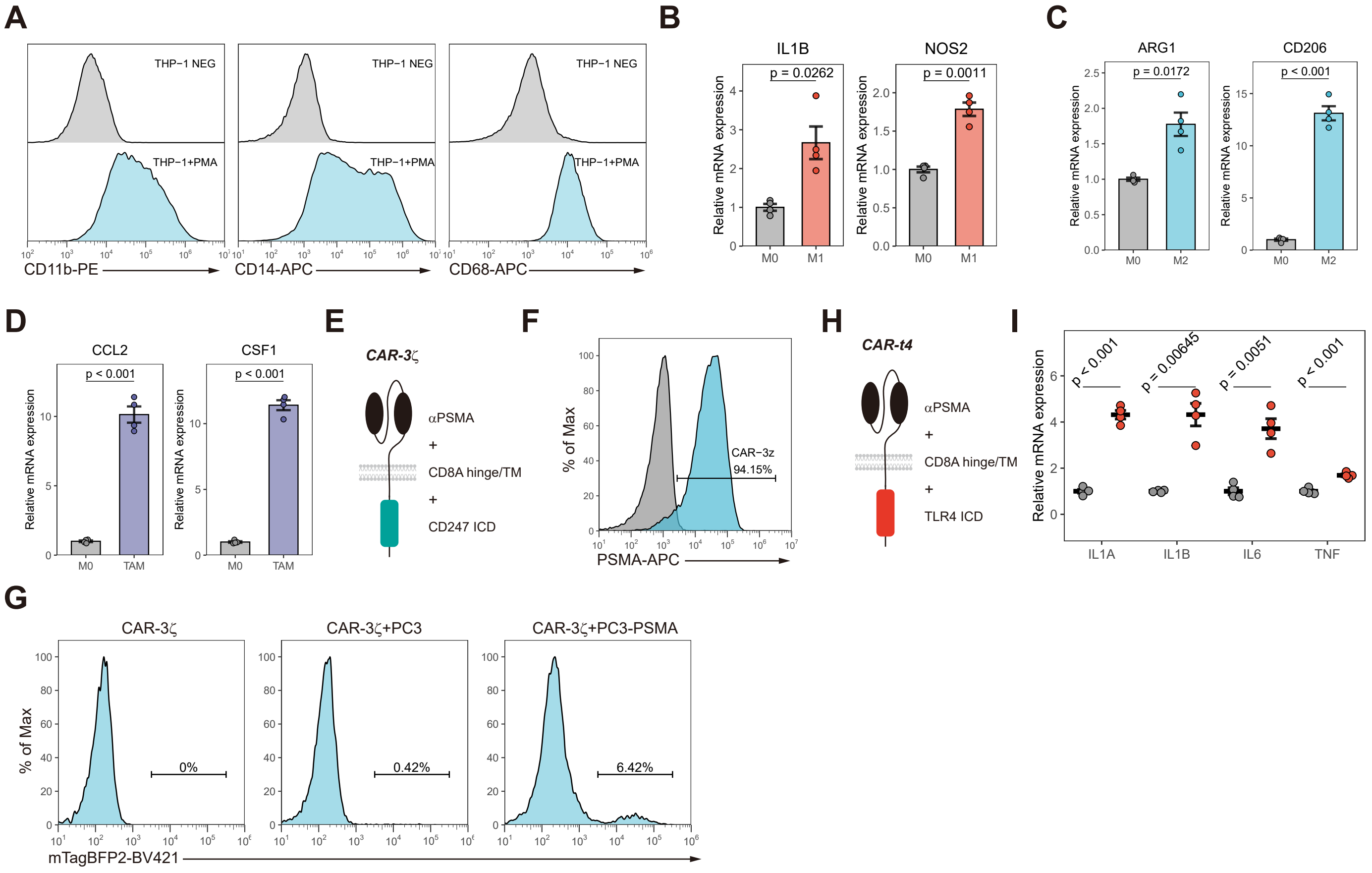


**Figure S1. THP-1-derived macrophages recapitulate the function of human primary macrophages in the context of CAR**

(A) Representative flow cytometric histograms of the surface expression levels of CD11b, CD14 and CD68 in THP-1 cells treated with PMA for 48 h.

(B-D) Relative mRNA expression levels of M1-, M2-, and TAM-associated genes in THP-1-derived macrophages (n = 4 biological replicates). M1 polarization was induced with LPS and IFNγ for 12 h. M2 polarization was induced using IL-4 for 48 h. TAM phenotype was generated by PC3 tumor cell conditioned media for 48 h.

(E) Schematic illustration of the CAR construct using CD247 intracellular domain (CAR-3ζ).

(F) Representative flow cytometric histograms showing CAR-3ζ surface expression.

(G) FACS-based phagocytosis of mTagBFP2+ PC3 or PC3-PSMA target tumor cells by CAR-3ζ macrophages.

(H) Schematic illustration of the CAR construct using TLR4 intracellular domain (CAR-t4).

(I) Relative mRNA expression of the M1-related genes in CAR-t4 macrophages treated with PSMA positive tumor cells for 24 h (n = 4 biological replicates).

The plots in (A, F, G) are representative of at least 3 independent experiments. Unless specified otherwise, data are presented as mean ± s.e.m. and analyzed using the two-tailed Student’s t-test.

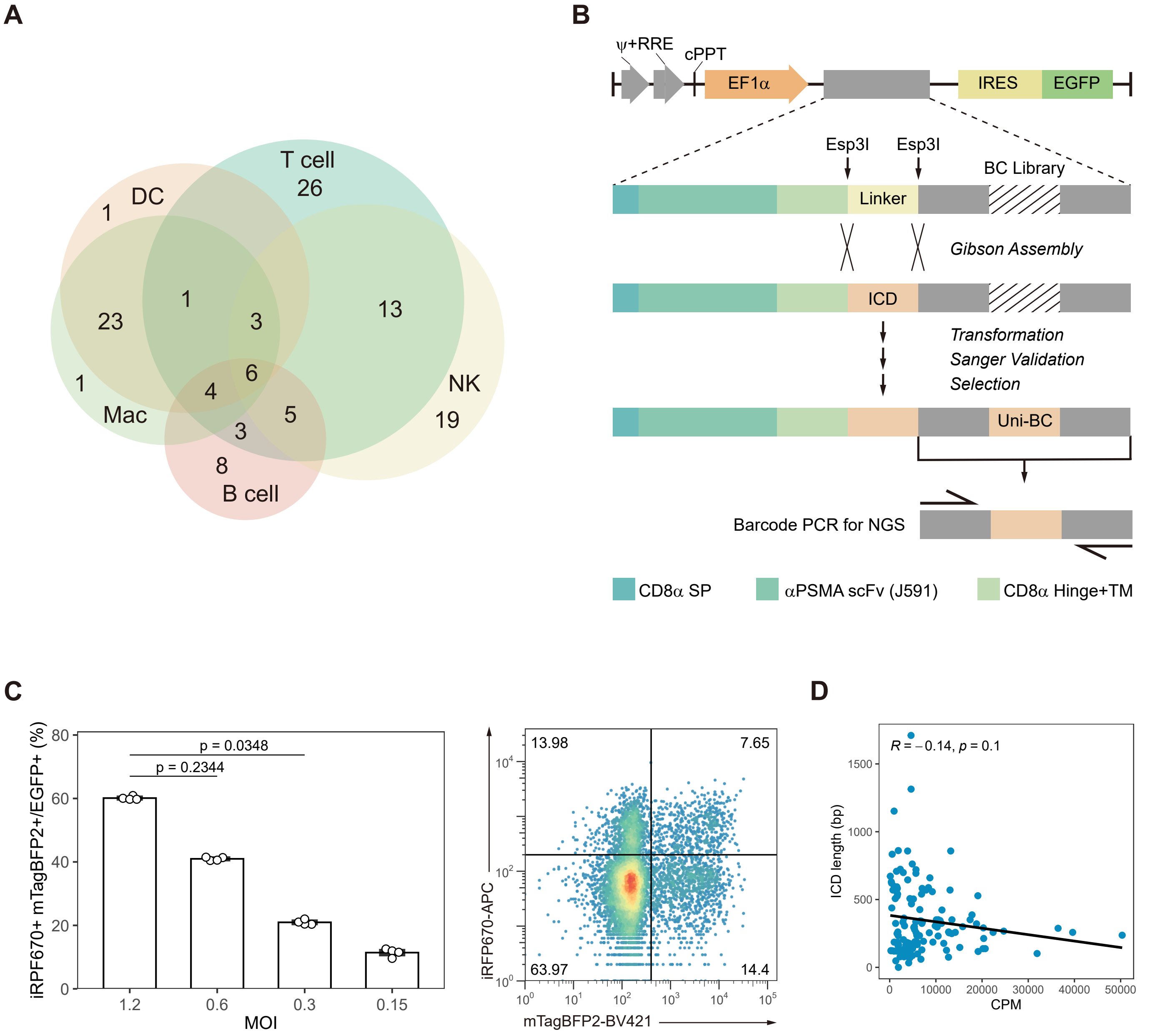


**Figure S2. Construction and validation of NaSDC CAR-M library**

(A) Venn diagram showing NaSDC library members across immune cell types. NK, natural killer cell. DC, dendritic cell. Mac, macrophage.

(B) Schematic representation of the generation of the NaSDC library. SP, signal peptide. TM, transmembrane. BC, barcode. Uni-BC, unique barcode. ICD, intracellular domain.

(C) Proportion of THP-1 cells expressing double positive (iRFP670$^+$ mTagBFP2$^+$) fluorescence (left). THP-1 cells were transduced with a mixed lentiviral pool containing iRFP670-EGFP and mTagBFP2-EGFP reporter lentiviruses at the specified MOI (n = 4). Representative flow cytometric plot showing iRFP670 and mTagBFP2 expression after transduction at MOI = 0.3 (right).

(D) Correlation of ICD lengths and their abundance in NaSDC THP-1 cells. CPM, counts per million.

Data in (C) are presented as mean ± s.e.m. and analyzed using one-way ANOVA followed by Dunnett's multiple-comparison test.

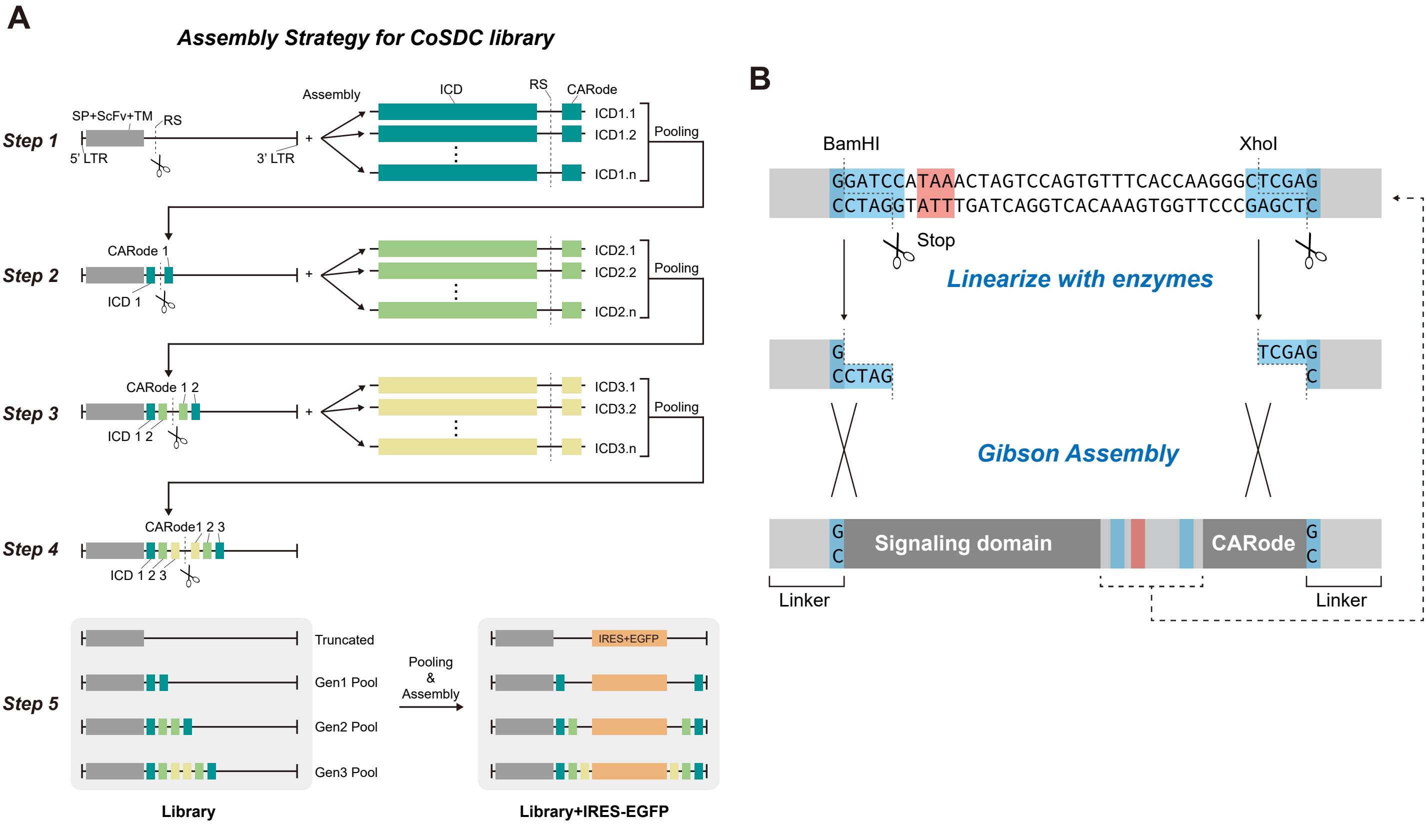


**Figure S3. Generation of CoSDC library using CARode method**

(A) Schematics describing the generation of CoSDC library. SP, signal peptide. TM, transmembrane. RS, restriction site. ICD, intracellular domain.

(B) CARode strategy for the combinatorial library construction.

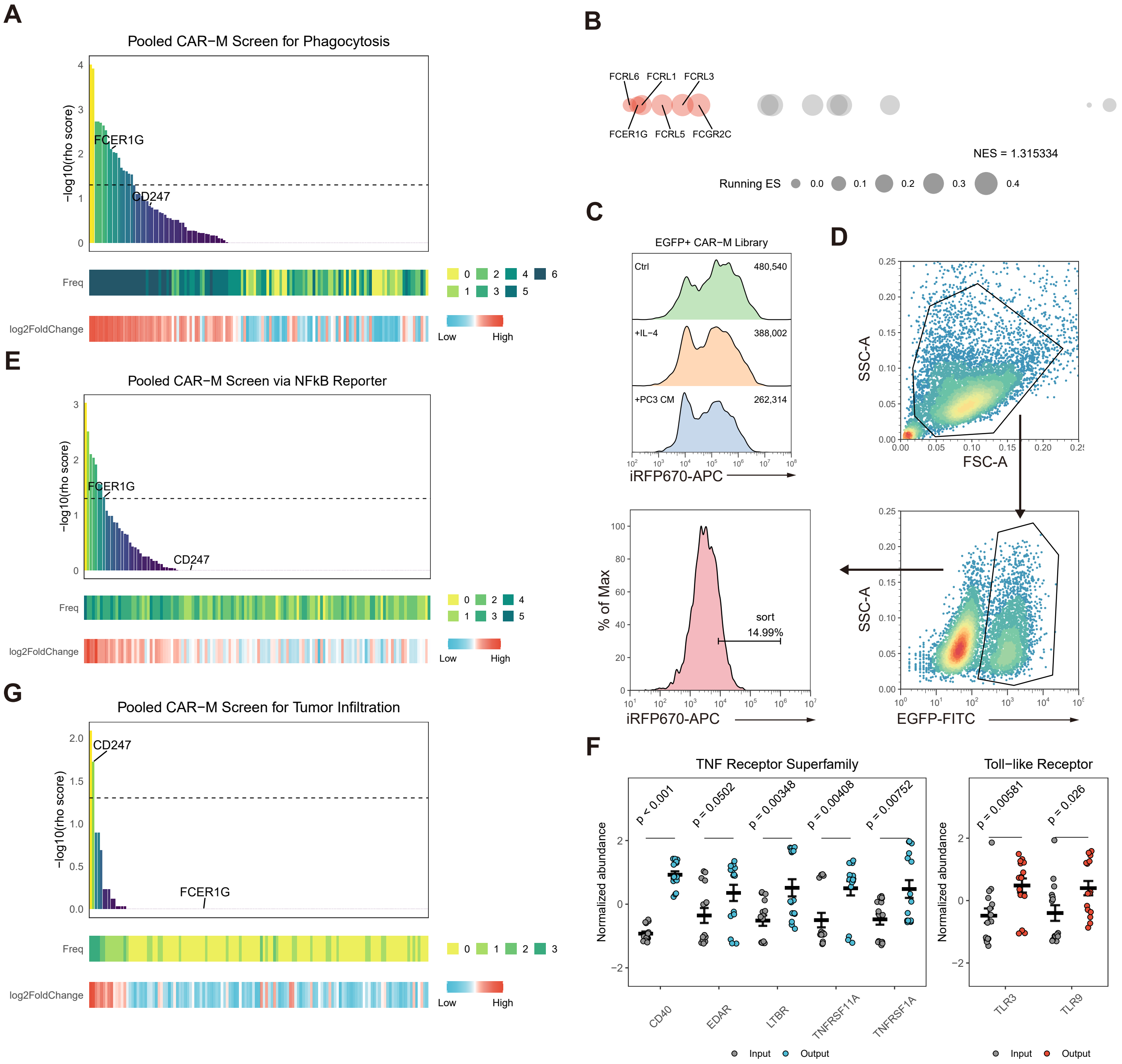


**Figure S4. NaSDC screens identify CAR-M variants with distinct anti-tumor functions**

(A) Bar plot of −log10 rho scores for all CAR variants in the pro-phagocytic screens (top). The dashed line represents −log10(0.05). Heatmap illustrating the frequency of each CAR variant identified in the upregulation list across six screening batches (middle). Heatmap showing the overall log2 fold change for each CAR variant in six batches of pro-phagocytic screens (bottom).

(B) Gene set enrichment analysis (GSEA) for Fc receptor family members used in pro-phagocytic screens. The red points represent the leading-edge CARs. NES, normalized enrichment score.

(C) Representative flow cytometric histograms of iRFP670 expression in the NFκB reporter NaSDC CAR-M library treated with IL-4 or PC3-conditioned media for 48 h. The plot is representative of at least 3 independent experiments.

(D) Gating strategy used for the pro-inflammatory CAR variant screens.

(E) Bar plot and heatmaps showing −log10 rho scores, frequency, and log2 fold change for pro-inflammatory CAR variant screens. The dashed line represents −log10(0.05).

(F) Z-score normalized abundance of the indicated CAR variants in the input and output pools of pro-inflammatory screens (n =15).

(G) Bar plot and heatmaps showing −log10 rho scores, frequency, and log2 fold change for pro-infiltrating CAR variant screens. The dashed line represents −log10(0.05).

Unless specified otherwise, data are presented as mean ± s.e.m. and analyzed using the two-tailed Student's t-test.

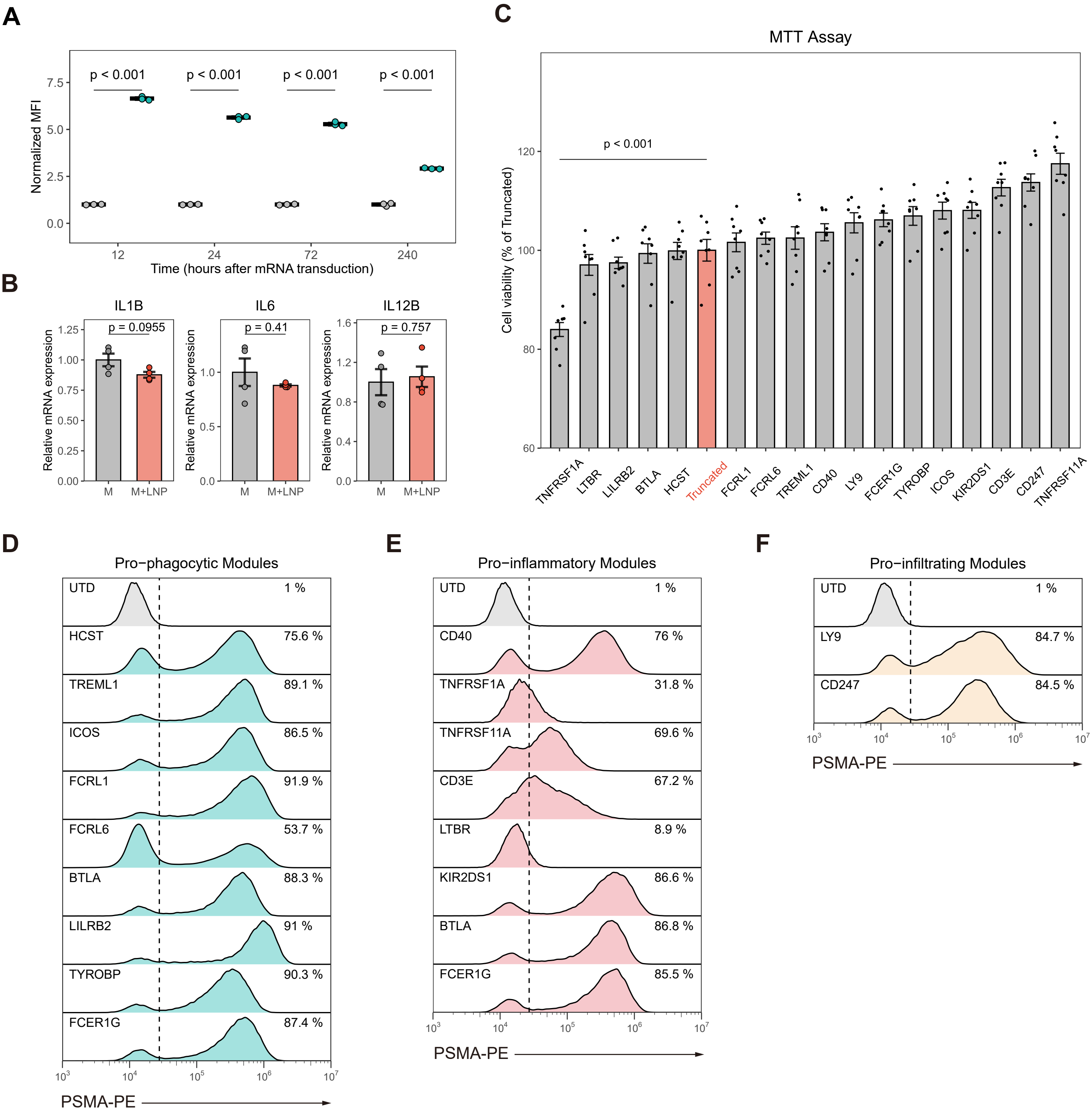


**Figure S5. Validation of CAR variants with distinct anti-tumor functions**

(A) MFI of hMDMs transduced with EGFP reporter mRNA. The EGFP expression was detected at the indicated timepoints. The flow cytometric signals are normalized by MFI of UTD macrophages (grey) (n = 3 biological replicates).

(B) Relative mRNA expression levels of M1-associated genes in hMDM treated with or without LNPs.

(C) Cell viability of hMDMs transduced with the specified CAR variants (n = 8 biological replicates).

(E-F) Flow cytometric histograms of surface expression levels of pro-phagocytic, pro-inflammatory, and pro-infiltrating CAR variants in hMDMs. Dashed lines represent the fluorescence intensity of the top 1% of UTD cells. The plots are representative of at least 3 independent experiments.

Unless specified otherwise, data are presented as mean ± s.e.m. and analyzed using the two-tailed Student's t-test.

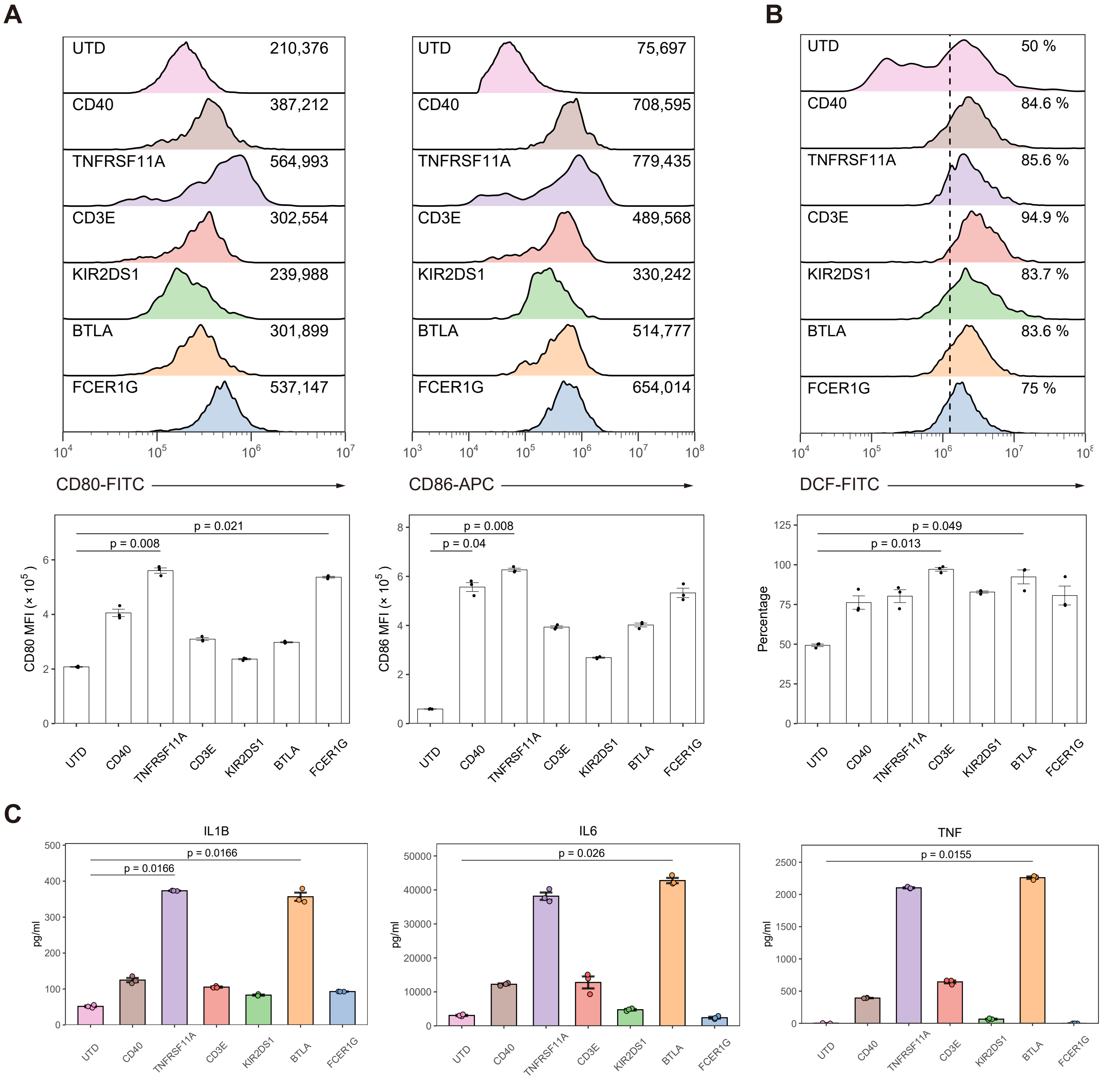


**Figure S6. NaSDC screens identify CAR-Ms with enhanced pro-inflammatory ability**

(A) Representative flow cytometric histograms and quantifications of surface levels of CD80 and CD86 in UTD or CAR-Ms.

(B) Representative flow cytometric histograms and quantifications of levels of ROS in UTD or CAR-Ms.

(C) IL-1β, IL-6 and TNFα secretion from the indicated CAR-Ms following 24 h coculture with antigen positive cells (n = 3 biological replicates). Protein levels were measured via ELISA.

The plots in (A, B) are representative of at least 3 independent experiments. Data in A-C are presented as mean ± s.e.m. and analyzed using one-way ANOVA followed by Dunnett's multiple-comparison.

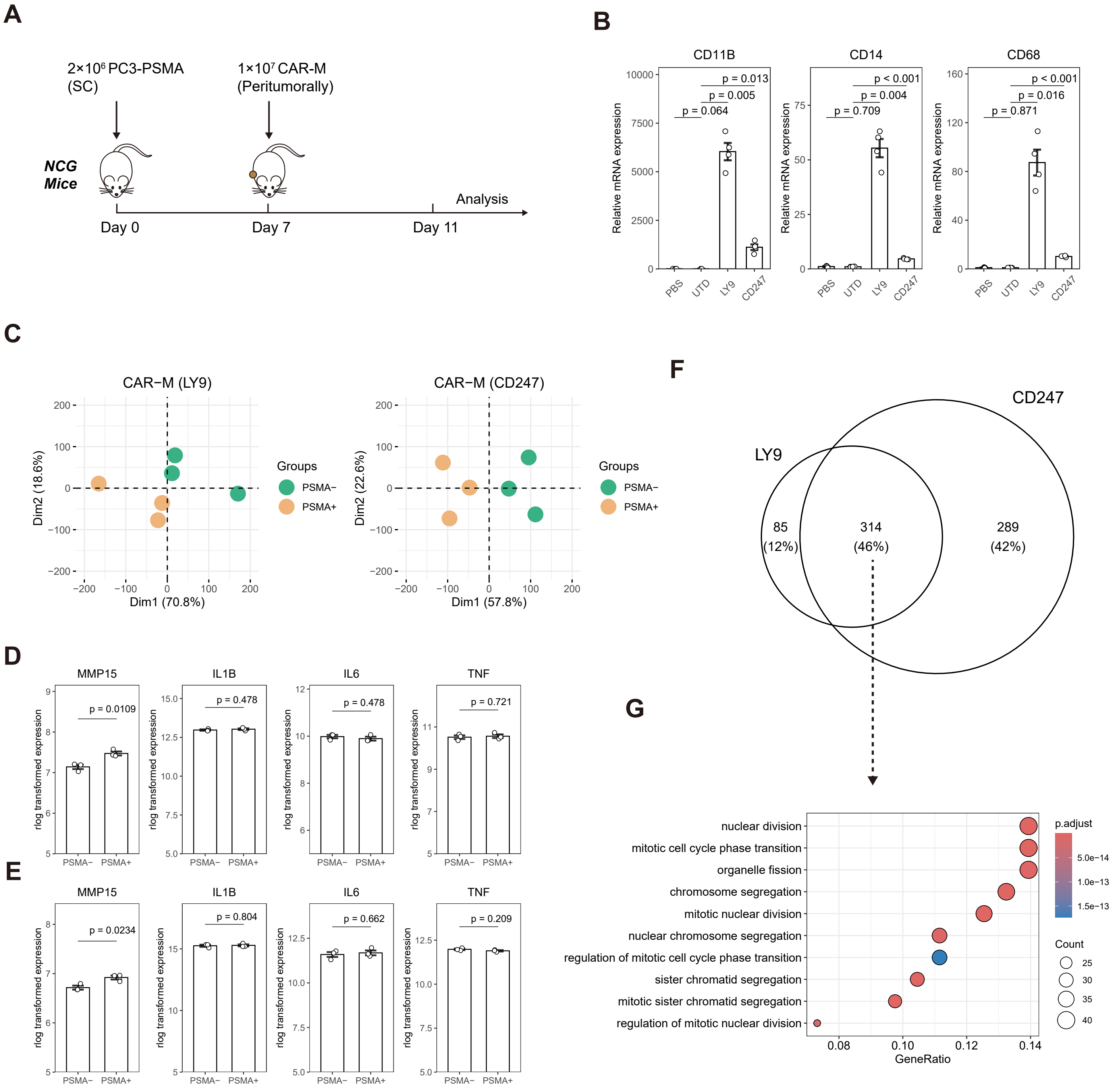


**Figure S7. NaSDC screens identify CAR-Ms with enhanced pro-infiltrating ability**

(A) Schematic illustration of experimental design. NCG mice were subcutaneously (SC) injected with $2 \times 10^6$ PC3-PSMA cells. Seven days after tumor administration, $1 \times 10^7$ UTD or CAR-Ms were injected peritumorally. Tumors were harvested four days after treatment.

(B) Relative mRNA expression of *CD11B*, *CD14* and *CD68* in tumor tissues treated with UTD or CAR-Ms (n = 4 biological replicates).

(C) Principal-component analysis (PCA) of RNA-seq data from CAR-Ms treated with K562 (PSMA-) cells or K562-PSMA (PSMA+) cells. Data were collected from 3 donors.

(D-E) The log transformed mRNA expression of the indicated genes in CAR-Ms (LY9 ICD) (D) and CAR-Ms (CD247 ICD) (E) treated with K562 cells or K562-PSMA cells (n = 3 donors).

(F) Venn diagram showing the relationships of upregulated genes between CAR-M (LY9) and CAR-M (CD247).

(G) GO enrichment analysis for the intersection genes.

Unless specified otherwise, data are presented as mean ± s.e.m. and analyzed using the two-tailed Student's t-test.

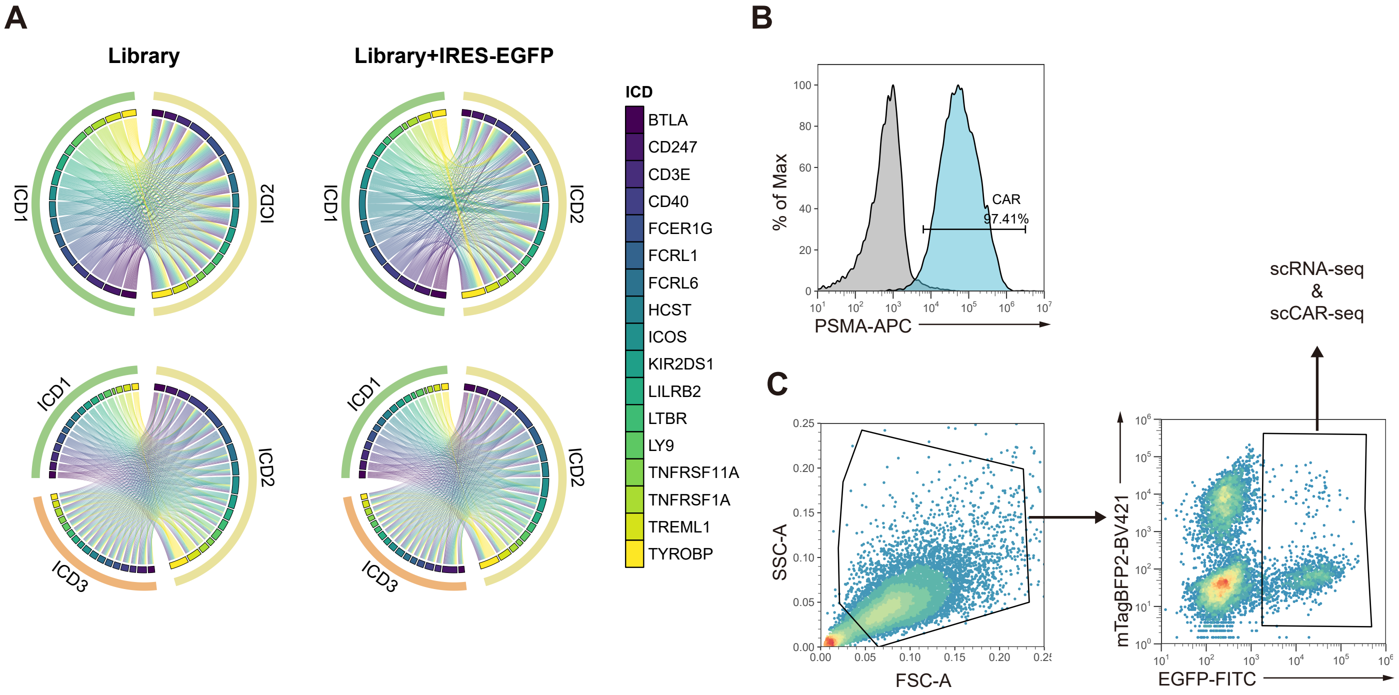


**Figure S8. Validation of CoSDC CAR-M library**

(A) Chord diagram showing the G2 and G3 CAR combinations that are present in CoSDC plasmid library with or without EGFP marker genes. The orders of ICDs are “CAR-ICD1-ICD2” or “CAR-ICD1-ICD2-ICD3”.

(B) Representative flow cytometric histograms of CAR expression in CoSDC library. Grey peaks represent the CAR expression levels in UTD. The plots are representative of at least 3 independent experiments.

(C) Gating strategy used for the scRNA-seq and scCAR-seq. Target tumor cells stably express mTagBFP2 fluorescent protein, CoSDC library cells express EGFP.

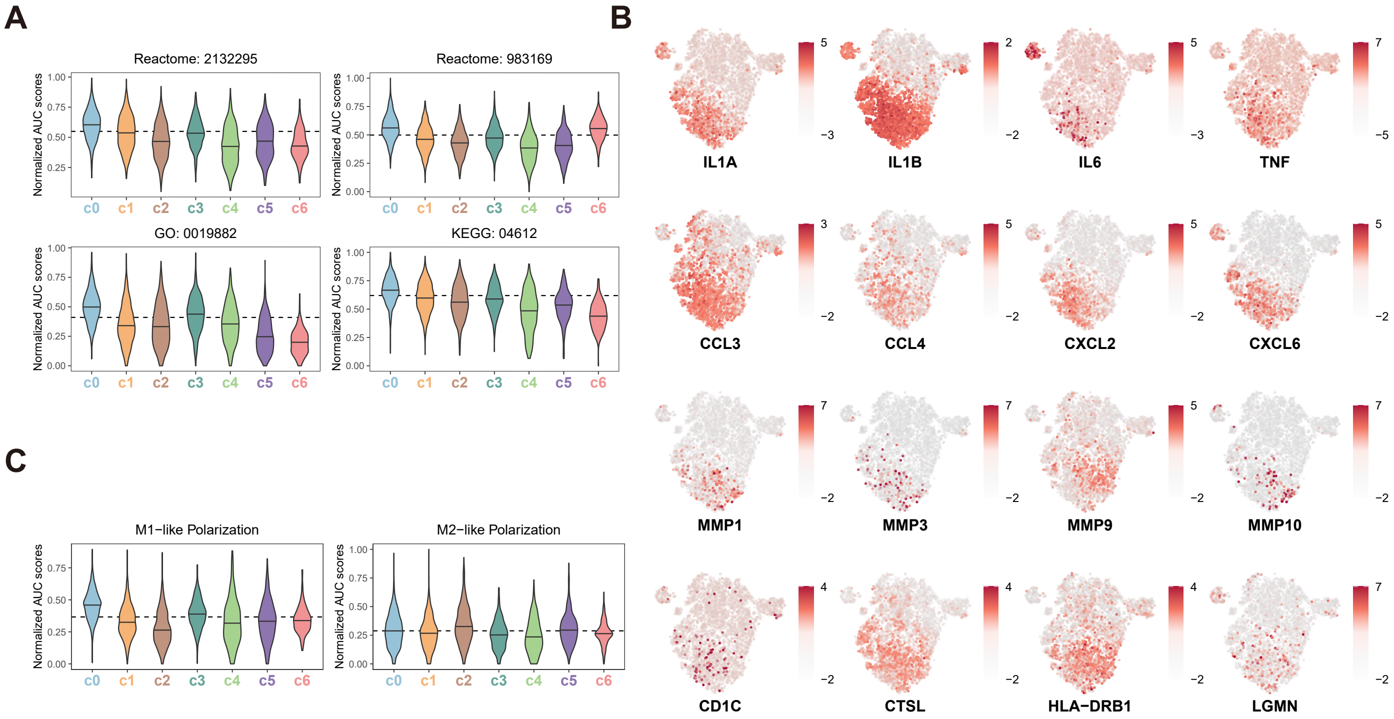


**Figure S9. scCAR-seq identify clusters with distinct transcriptional features**

(A) Normalized AUC scores for antigen processing and presentation-related gene sets. Reactome gene sets “MHC class II antigen presentation” (2132295) and “Class I MHC mediated antigen processing & presentation” (983169), GO gene sets “Antigen processing and presentation” (0019882), and KEGG pathway “Antigen processing and presentation” (04612) were used. Dashed lines represent the AUC scores of CARΔ.

(B) Feature plots showing the expression levels of the indicated genes.

(C) Normalized AUC scores for M1 activation and M2 activation gene sets. Dashed lines represent the AUC scores of CARΔ.

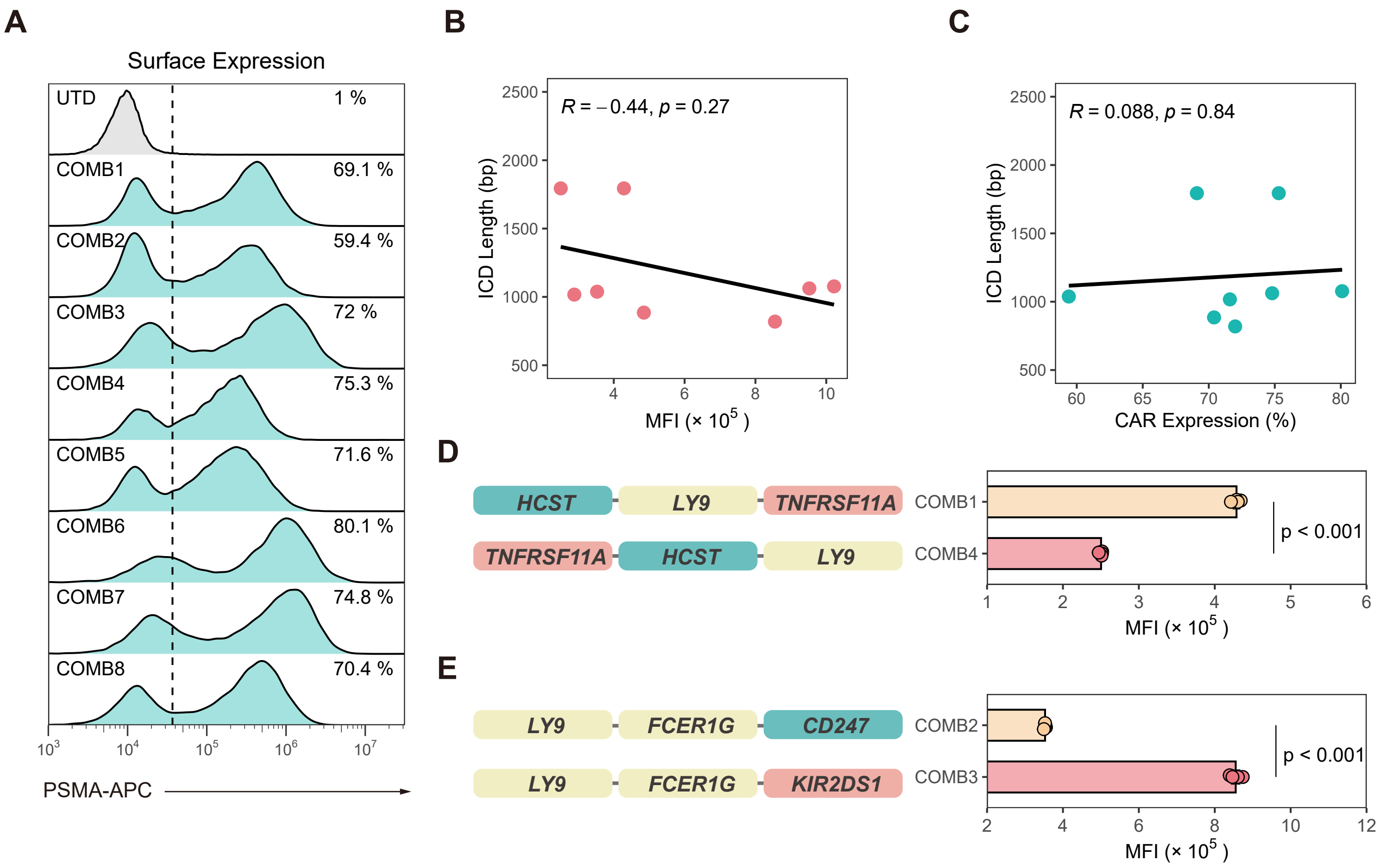


**Figure S10. ICD types and arrangements are correlated with CAR expression levels**

(A) Flow cytometric histograms of CAR expression for the indicated CAR-M variants. The plots are representative of at least 3 independent experiments. The dashed line represents the fluorescence intensity of the top 1% of UTD cells.

(B) Correlation between ICD lengths and their expression levels (MFI) in hMDMs.

(C) Correlation between ICD lengths and their positive rates (%) in hMDMs.

(D) MFI quantification of CAR expression in COMB1 and COMB4 CAR-Ms (n = 4).

(E) MFI quantification of CAR expression in COMB2 and COMB3 CAR-Ms (n = 4).

Unless specified otherwise, data are presented as mean ± s.e.m. and analyzed using the two-tailed Student's t-test.

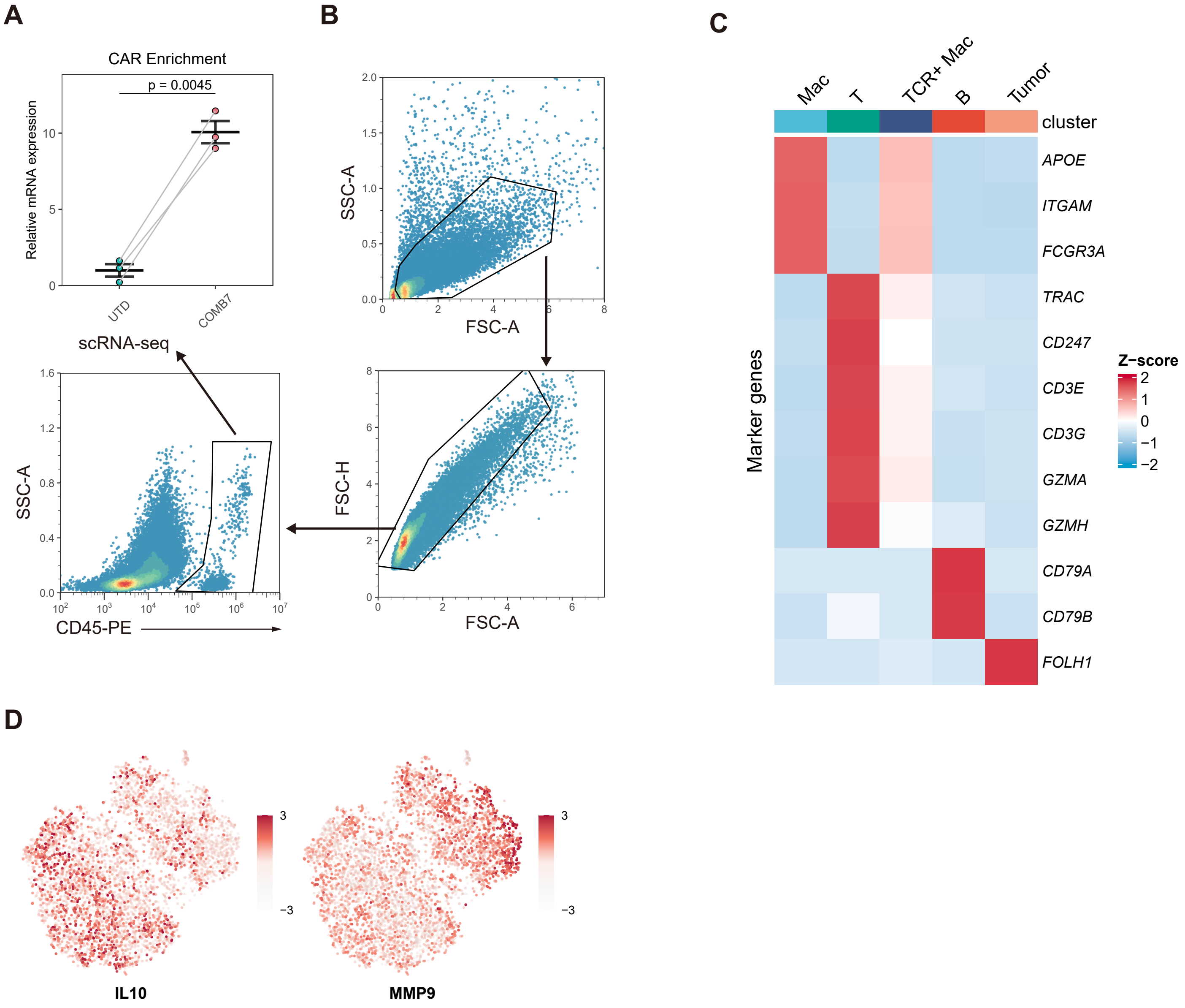


**Figure S11. scRNA-seq identifies TME of humanized immune system mouse model**

(A) Relative mRNA expression of CAR-encoded genes in matched tumor tissues (n = 3 mice).

(B) Gating strategy used for the scRNA-seq.

(C) Heatmap showing the expression levels of marker genes used in cell cluster identification. Mac, macrophage. T, T cell. B, B cell.

(D) Feature plots showing the expression levels of the indicated genes.

Data in (A) are presented as mean ± s.e.m. and analyzed using the paired t-test.

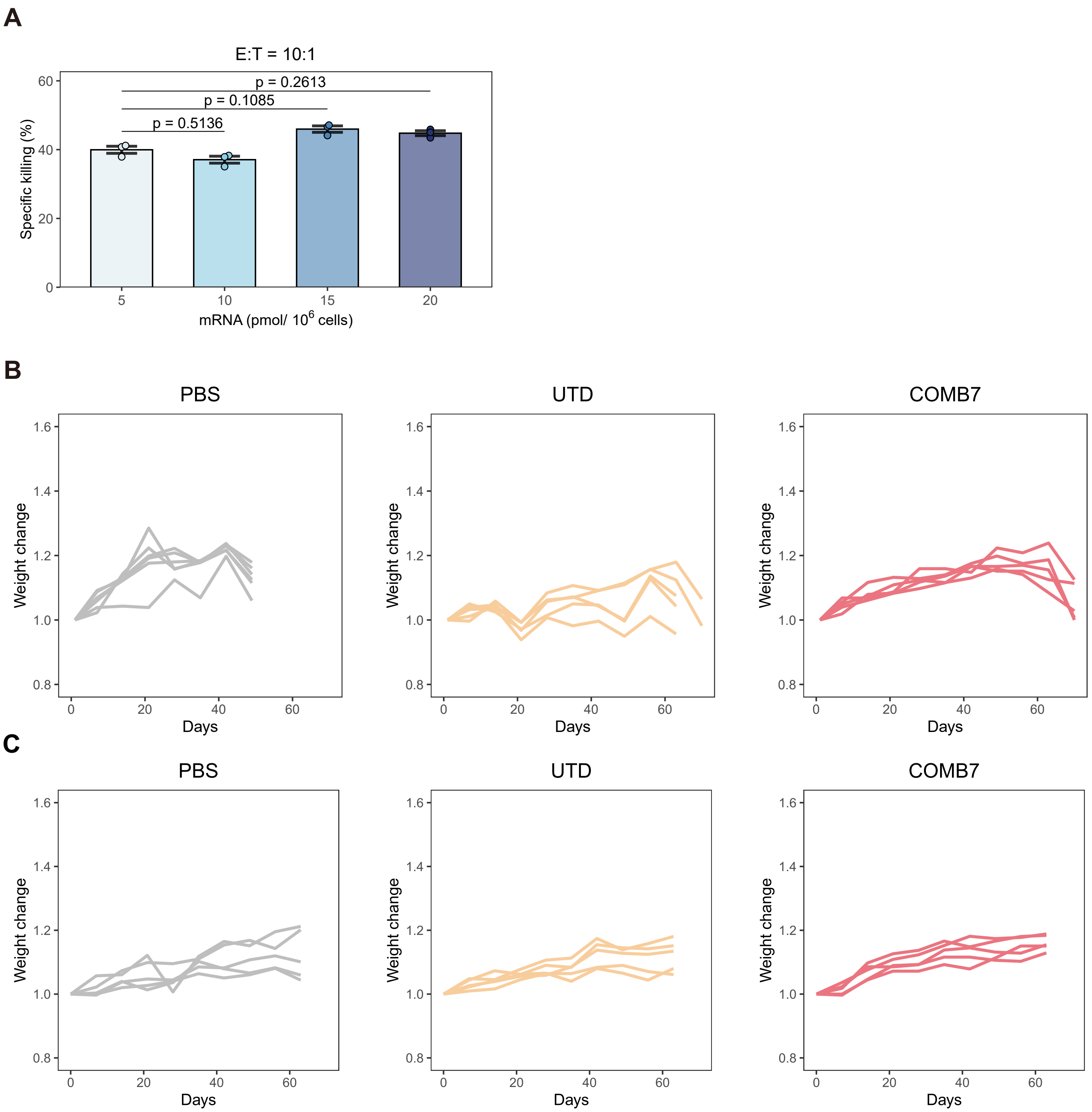


**Figure S12. Validation of CD40-LY9-FCRL1 CAR-Ms in vitro and in vivo**

(A) Specific killing of PC3-PSMA target cells by hMDMs (E/T ratio = 3/1) at the indicated COMB7 CAR mRNA transduction conditions (n = 3).

(B-C) Weight loss following tumor IP (B) or SC (C) injection was monitored routinely. Data shown in the plots are individual values.

Data in (A) are presented as mean ± s.e.m. and analyzed using one-way ANOVA followed by Dunnett's multiple-comparison.
